# isoGS: A genomic method to navigate the breeder’s dilemma of diversity versus eliteness

**DOI:** 10.64898/2026.11.23.768689

**Authors:** Theodore Monyak, Safiétou Tooli Fall, Kenneth Opare-Obuobi, Manuel Q. Esguerra, Allan Fritz, Clara Cruet-Burgos, Geoffrey P. Morris

## Abstract

Global networks of public crop breeding programs exchange germplasm to share traits and increase diversity for genetic gain. There is a fundamental dilemma in crop breeding between the need to expand the genetic variation in elite gene pools while avoiding disruptions to essential traits required for eliteness. One particular challenge in admixed germplasm is the release of undesired transgressive segregation due to the disruption of favorable epistatic haplotypes of essential traits, which may stymie deployment of genomic selection (GS) In this study, we used breeding simulations to model recurrent population improvement for an annual crop species that can be selfed and crossed to examine the effectiveness of GS in admixed germplasm. We quantified the genetic heterogeneity between founder lines with an “isoeliteness” score, an effect size-weighted Hamming distance using quantitative trait loci markers. Populations with low founder isoeliteness (i.e. low genetic compatibility) had lower genetic gain with GS in early cycles than those with high founder isoeliteness (63% and 27% lower after 2 and 4 cycles). To mitigate this, we developed an isoeliteness-guided GS (isoGS) method to recover favorable epistatic haplotypes in a segregating population, which enabled 31% and 15% more genetic gain after 4 and 20 cycles. Our results suggest that founder isoeliteness scores and isoGS are valuable tools for overcoming the diversity vs. eliteness dilemma by managing the introduction of genetic diversity in a trait-specific manner.

**ARTICLE SUMMARY:** A fundamental dilemma in breeding exists between maintaining genetic diversity that enables genetic gain and minimizing variation underlying essential traits that constitute suitability to local preferences. Global networks of public crop breeding programs exchange germplasm to share traits and increase genetic diversity, but lack strategies for managing genetic heterogeneity, which can disrupt the stability of essential traits and lead to wasted breeding resources. We developed a framework called “isoGS” that directly accounts for hidden allelic interactions, and used breeding simulations to demonstrate how it can help breeders manage a wide base of genetic variation while minimizing detrimental short-term breeding inefficiencies.

## INTRODUCTION

Crop improvement depends on the continued availability of genetic variation as the raw material for selection. Yet while genetic diversity is beneficial to drive genetic gain for polygenic *desired* traits such as yield (Cobb, Juma, et al. 2019; Rutkoski 2019b; Mackay et al. 2021), it can be detrimental for essential *attained* traits associated with eliteness, such as flowering time and plant height (Fall et al. 2026). This tradeoff between diversity and eliteness is not captured by the breeder’s equation, Δ*G = (σ_a_ * r * i)/L*, which only shows how the expected genetic gain Δ*G* depends on additive genetic variance (*σ_a_)*, selection accuracy (*r*), selection intensity (*i*), and cycle length (*L*) (Lush 1937; Falconer and Mackay 2009: 189). However, this tradeoff underscores a core dilemma in breeding: that in order to minimize disruption to attained traits, breeding programs favor crosses between highly related elite materials, thus narrowing the genetic bases of their germplasm and limiting the potential for genetic gain (Kannenberg and Falk 1995; Cooper et al. 2001; Swarup et al. 2021). Standard pre-breeding (Simmonds 1993) can address this dilemma by enabling the introduction of novel genetic variation, but this is an expensive option beyond the means of many under-resourced breeding programs. In contrast with the closed breeding paradigm, for certain crops and geographies, networks of public sector breeding programs routinely exchange germplasm and information due to historical legacy, resource constraints, and an aim to increase their collective capacity to respond to new agricultural constraints (Atlin et al. 2017; Muleta et al. 2022; Das et al. 2024). It is critical that this germplasm exchange is accompanied by strategies to minimize disruption for eliteness in the resulting admixed populations by managing genetic diversity in a trait-specific manner.

Genetic heterogeneity, the presence of allelic variation underlying traits of interest, is a particular challenge with attained traits: essential phenotypes that play a large role determining the fitness of a genotype. Though phenotypes may be common across breeding programs, different programs may have distinct alleles obtained from intensive stabilizing selection for each trait, due to genotypic degeneracy (Edelman and Gally 2001; Barghi et al. 2020). When phenotypically similar but genetically heterogeneous *allo-*elite lines (Fall et al. 2026) are crossed, a high proportion of the progeny can be transgressive segregants (Rieseberg et al. 1999; Paaby and Rockman 2014) that must be discarded. Conversely, if *iso-*elite parents (isogenic for alleles at quantitative trait loci (QTL) underlying attained traits) are crossed, their progeny are expected to largely retain their elite attained trait phenotypes. For example, U.S. great plains winter wheat breeding programs are typically fixed for one of the *Rht1* or *Rht2* semi-dwarfing alleles (Allan 1989), and try to avoid segregation at these loci caused by allo-elite crosses. In the case of the various mutant alleles of the six classical maturity genes underlying flowering time in global sorghum (Quinby 1967; Rooney and Aydin 1999), this challenge is exacerbated by epistasis (Yang et al. 2014; Casto et al. 2019).

Epistasis among the oligogenic components of attained traits is a challenge for breeding efforts in heterogeneous germplasm, particularly for programs using or considering genomic selection (GS) (Meuwissen et al. 2001; Heffner et al. 2009). Standard GS methods presume that phenotypic variation in quantitative traits is underlied by many additive QTL of small effect (Fisher 1919; Podlich and Cooper 1999; Barton et al. 2017; Morgante et al. 2018). However, epistasis is not only a frequent observation in global pre-breeding resources (Yu et al. 1997; Brown et al. 2011; Li et al. 2015; Olatoye et al. 2020), but an emergent property at the fitness level for any oligogenic additive trait under stabilizing selection (Wright 1935; Barton 1986; Whitlock et al. 1995). When a phenotype is determined by intergenic interactions, in the case of biological epistasis, or offsetting allelic effects in the case of fitness epistasis, the combination of alleles fixed in a population can be considered a co-adapted haplotype complex (Bernardo 2009; Condón et al. 2009; Volis et al. 2011). Allo-elite crosses can disrupt these haplotypes and release unwanted epistatic variance for attained traits (Cooper et al. 2005; Mackay 2014; Monyak and Morris 2026), which can hinder direct selection for additive variation (Technow et al. 2021).

Genetic characterization of breeding lines may assist with minimizing undesirable epistatic variance. Genomic markers may be used for QTL-based genotypic inference (Fall et al. 2026) to identify genetically compatible breeding lines, while marker-assisted selection (Bouchez et al. 2002; Cobb, Biswas, et al. 2019) may be used to prioritize the preservation of favorable co-adapted epistatic haplotypes underlying attained traits. In this study, we used breeding simulations to investigate genomic marker-based approaches of mitigating fitness epistasis in heterogeneous germplasm. Our simulations modeled recurrent population improvement in an annual crop species that can be selfed and crossed. We demonstrated the importance of founder genetic compatibility for genomewide prediction (GWP) accuracy, genetic gain, and cost-effectiveness, and developed an isoeliteness-guided GS framework based on elite haplotypes. These methods can help breeders manage a wide base of genetic variation while minimizing detrimental short-term breeding inefficiencies.

## MATERIAL AND METHODS

### Simulation of a genotype-to-fitness map

The genotype-to-fitness map used in this study was the same as was described in (Monyak and Morris 2026) (see Table 1). Briefly, a founder population of 2000 diploid organisms with 10 chromosomes was simulated using coalescent MaCS software (Chen et al. 2009) with the R package “AlphaSimR” (Gaynor et al. 2021). There were two additive oligogenic attained traits (*t_1_, t_2_)* (e.g. flowering time, plant height) with *L* = 10 QTL each, representative of traits with an optimal phenotype under stabilizing selection. There was one additive polygenic desired trait (e.g. yield potential) (*t_3_)* with *L* = 500, representative of a trait under directional selection. There was a geometric distribution of allele substitution effect sizes for the oligogenic traits, in which the relative decrease in effect size for each subsequent QTL, *a,* was determined based on *L* as: *a* = (*L* − 1) / (*L* + 1) (Lande and Thompson 1990). The allele substitution effect sizes for the polygenic trait were randomly drawn from a gamma distribution.

**Table 1.**
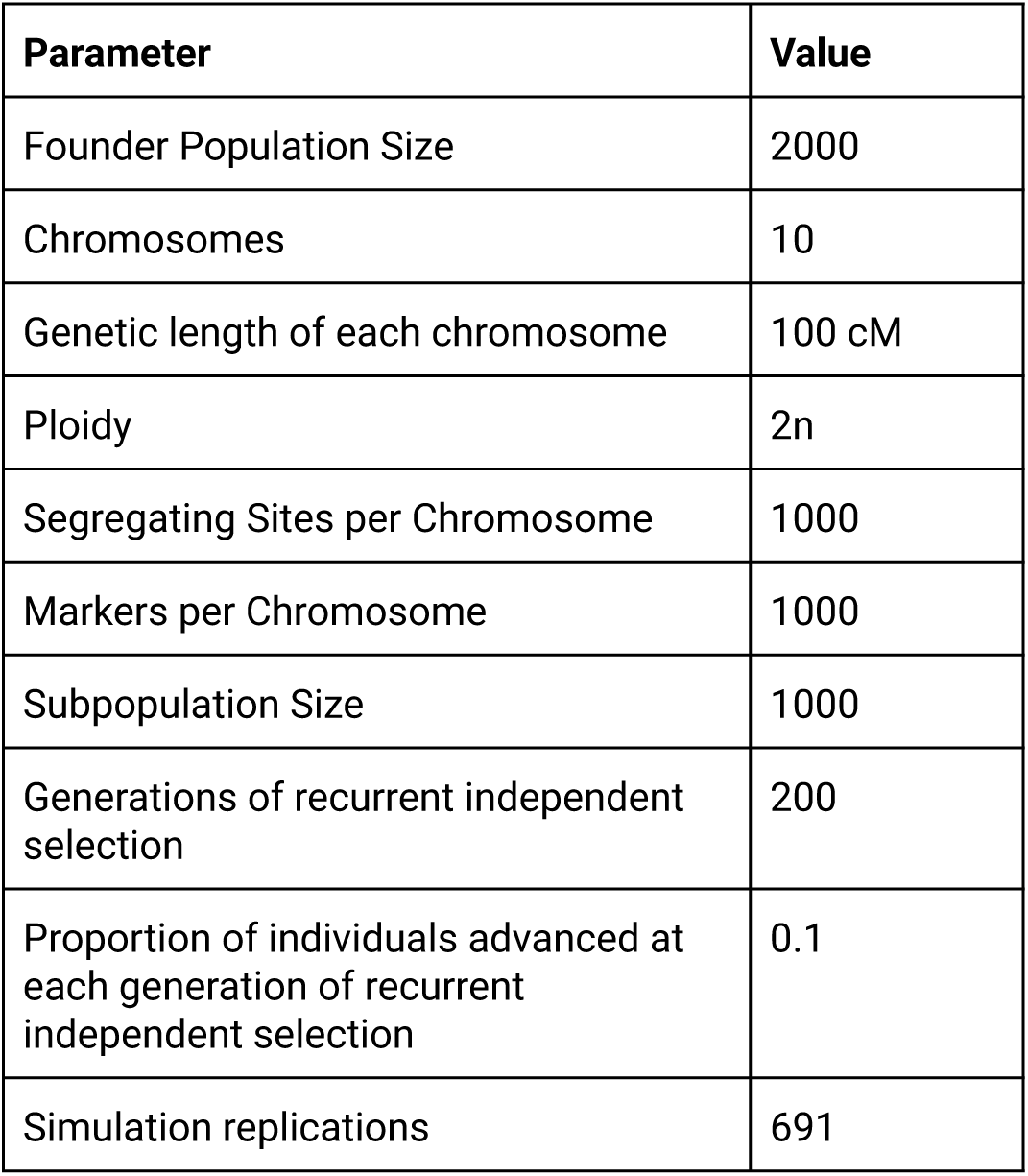
Population establishment parameters.

The adaptedness of a genotype to an environment was quantified with a ‘suitability’ (*S)* score, based on the distance of each of the attained trait phenotypes from an optimal phenotype of ‘0’. *S* was calculated as 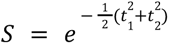, producing a score between 0 and 1 (optimal). ‘Realized yield’ (i.e. fitness) (*W*) was calculated as *S* * *t*_3_, effectively penalizing yield potential by the maladaptedness of an individual. Though *W* was a simplified composite trait lacking in biological realism, it was a useful model for investigating emergent fitness epistasis that can affect traits with an additive genetic architecture under stabilizing selection. Narrow-sense heritability (*h^2^*) was initially set to 0.1 for the oligogenic attained traits and 0.05 for the polygenic desired trait, representative of the low heritability expected in landrace cultivation.

### Development of RIL families

A simulation pipeline (Table 2) was developed to represent the development of two independent breeding programs, followed by germplasm exchange between those programs (Fig. 2a). Results from 2000 unique replications were analyzed. Initially, 2 subpopulations of 1000 individuals were randomly sampled from the founder population, and underwent independent recurrent selection for 200 generations. At each generation, the top 10% of individuals in each subpopulation ranked by realized yield were advanced and randomly mated to produce the next generation. These 200 generations of selection represented the independent development of landraces, with a sufficient number of generations for each subpopulation to approach fixation for both attained traits. Following this stage, two inbred lines (i.e. purified landraces) were developed from each subpopulation by intensively selecting each subpopulation for *‘S’*. This was representative of the process in which nascent breeding programs turned heterogeneous landrace materials into stable founders for their germplasm. To reflect the reduced environmental variance in a controlled breeding scenario, *h^2^* was increased to 0.8 for the attained traits and 0.3 for the desired trait.

**Table 2.** Trait parameters.

| **Parameter** | **Traits 1 and 2** | **Trait 3** |
|---|---|---|
| QTL per chromosome | 1, 2, or 5 | 50 |
| Narrow-sense heritability | 0.1, then 0.8 | 0.05, then 0.3 |
| Initial mean genetic value | 2 | 50 |
| Initial genetic variance | 0.4 | 50 |
| Effect size sampling distribution | Custom geometric distribution | Normal |

To model a genetically heterogeneous open breeding population, an admixed RIL family was developed by crossing purelines derived from each of the two subpopulations. These purelines were assumed to be allo-elite. The initial cross produced 250 F_1_ individuals, which were selfed to produce 4 F_2_ progeny per individual. Each F_2_ was selfed repeatedly through single-seed descent to generate an F_10_ recombinant inbred line (RIL) family with 1000 lines. This admixed RIL family was contrasted with an unadmixed RIL family, derived instead from a biparental cross of the two purelines developed from the same subpopulation, assumed to be iso-elite. The unadmixed family was representative of a genetically homogeneous closed breeding program germplasm.

### Measuring founder isoeliteness

To quantify the genetic heterogeneity at the QTL underlying a trait, the *isoeliteness* between each pair of RIL founders was calculated as a score between 0 (contrasting alleles at all QTL) and 1 (isogenic at all QTL) (Monyak and Morris 2026). Since all the selected lines were adapted, a lower isoeliteness score therefore indicated that two lines were more allo-elite. An isoeliteness score of 1 indicated that all QTL were isogenic and a score of 0 reflected contrasting alleles at all QTL. Isoeliteness between individuals *a* and *b* was calculated as:

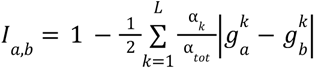

Where 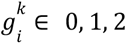 was the allelic dosage at QTL *k* for individual *i* (2 = homozygous for allele 1, 0 = homozygous for allele 0 and 1 = heterozygous), and the weight term *α_k_* / *α_tot_* was the allelic substitution effect at QTL *k* over the sum of allelic substitution effects across all QTL for that trait. RIL families were categorized based on the mean isoeliteness across the two attained traits with respect to the initial founders as “low” (mean isoeliteness ≤ 0.8), “moderate” (mean isoeliteness between 0.8 and 0.95), and “high” (mean isoeliteness ≥ 0.95). Additionally, the “desired trait heterogeneity” was calculated using the isoeliteness formula, using the QTL underlying *t*_3_.

### Enriching favorable haplotypes underlying isoeliteness

To enrich haplotypes underlying eliteness in a segregating population derived from a RIL family, a second isoeliteness-based framework was developed: isoeliteness-guided genomic selection (isoGS) (Fig. 1). When used with phenotypic selection (PS), this approach was referred to as isoPS. This approach was based on the assumption that each subpopulation contained a set of distinct co-adapted epistatic complexes that were disrupted in the admixture through recombination. Since all traits had a purely additive genotype-to-trait mapping, any detected epistasis would be due to the emergence of fitness epistasis for traits under stabilizing selection. Epistatic linkage mapping was run with the R package “R/qtl” (Arends et al. 2010) on the RIL family with a high-density SNP array of 100 markers per chromosome for the realized yield phenotype. Then, each individual in an F_2_ population derived from this RIL family was genotyped at the markers colocalized with each significant detected interaction peak.

**Figure 1.**
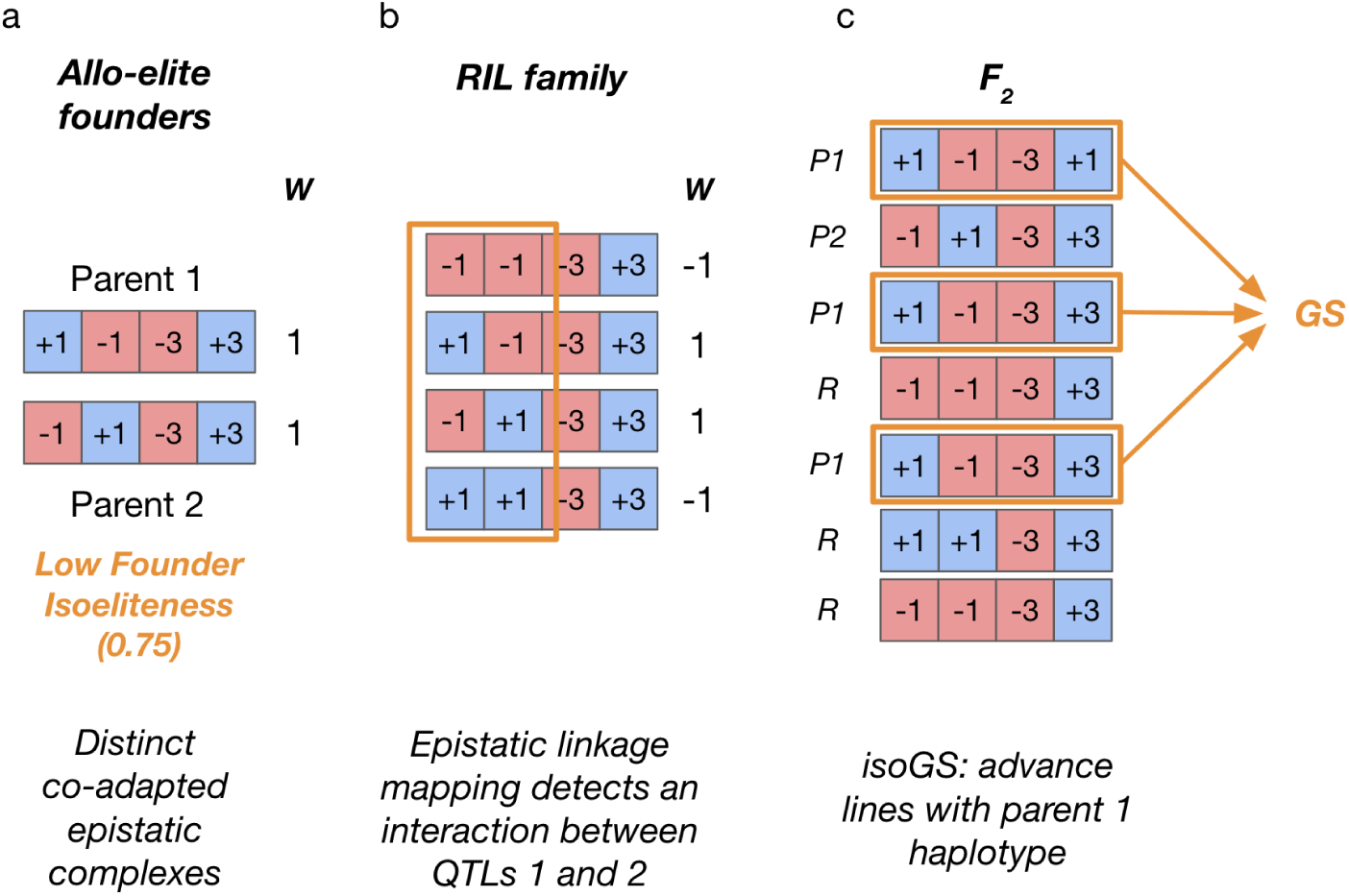
isoGS schematic diagram. In this illustrative example, there is only one trait with four QTL, with additive effect sizes indicated in each QTL square. A simplified fitness function is used here, where fitness (*W)* is 1 (optimal) if the trait phenotype is 0, and −1 (suboptimal) otherwise. Two subpopulations are assumed to have evolved unique co-adapted epistatic complexes through independent recurrent selection. (a) Pureline allo*-*elite parents derived from each subpopulation have contrasting haplotypes at the epistatic loci, though they both have optimal fitness, and they have an isoeliteness score of 0.75. (b) The epistatic complexes then become disrupted through recombination when the parents are crossed, leading to segregation in the resulting admixed RIL family. Epistatic linkage mapping detects a significant interaction with respect to *W* between the two segregating QTL. (c) An F_2_ population derived from intercrossed RILs is genotyped, and each individual is categorized as having a “parent 1”, “parent 2” or “recombinant” haplotype across the epistatic QTL. Since there are more individuals with the “parent 1” haplotype than with the “parent 2” haplotype, only those with the “parent 1” haplotype are selected to be advanced.

For each interaction, each line was characterized as having the ‘parent 1 (P1)’, ‘parent 2 (P2)’, or ‘recombinant (R)’ haplotype by comparing its haplotype at those two markers to that of the RIL parents. For each interaction, the “majority haplotype” in the F_2_ population was determined, and each individual’s haplotype was coded as a ‘1’ if it matched that majority, and a ‘0’ otherwise. Next, the individuals were hierarchically ranked by the number of ‘1’s’ they possessed across each interaction (from highest LOD score to lowest), and lastly by realized yield. Then, the top 25% of the ranked individuals, representing those individuals with the most “majority parental haplotypes” (i.e. a co-adapted epistatic combination of alleles) were selected and advanced. In addition to using the high-density SNP array, isoGS was performed in parallel using two alternative sets of markers: a low-density SNP array with 20 markers per chromosome: ‘isoGS (Low Density)’, and the actual QTL underlying the attained traits: ‘isoGS (Perfect Markers)’.

### Evaluation of genomewide prediction accuracy

The GWP model used in simulations was ridge regression best linear unbiased prediction (RR-BLUP) (Meuwissen et al. 2001). The RR-BLUP model was fitted to realized yield phenotypes using the RRBLUP() function in AlphaSimR (Gaynor et al. 2021). The RR-BLUP procedure fits the following model:

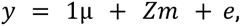

where *y* is a vector of phenotypes, *μ* is an vector of the overall mean, *Z* is the marker incidence matrix*, m* is the vector of normally distributed marker effects (n = 100 markers per chromosome), and *e* is a normally distributed vector of residuals. The GWP model accuracy (*r*_MG_) in a target population was the Pearson correlation between the model’s genomic estimated breeding values (GEBVs) and the genetic values calculated from *t_1_, t_2_, and t_3_*, calculated using the cor() function in base R (R Core Team 2025). GWP accuracy in RIL families was evaluated in three different scenarios (Fig. 2a). For the “within-family” scenario, 5-fold cross validation accuracy was calculated within each unadmixed RIL family. For the “cross-family” scenario, the training population (TP) was the unadmixed RIL family derived from one subpopulation, and the target population was the unadmixed RIL family derived from the other subpopulation. For the “admixed” scenario, 5-fold cross-validation accuracy was calculated within the admixed RIL family.

**Figure 2.**
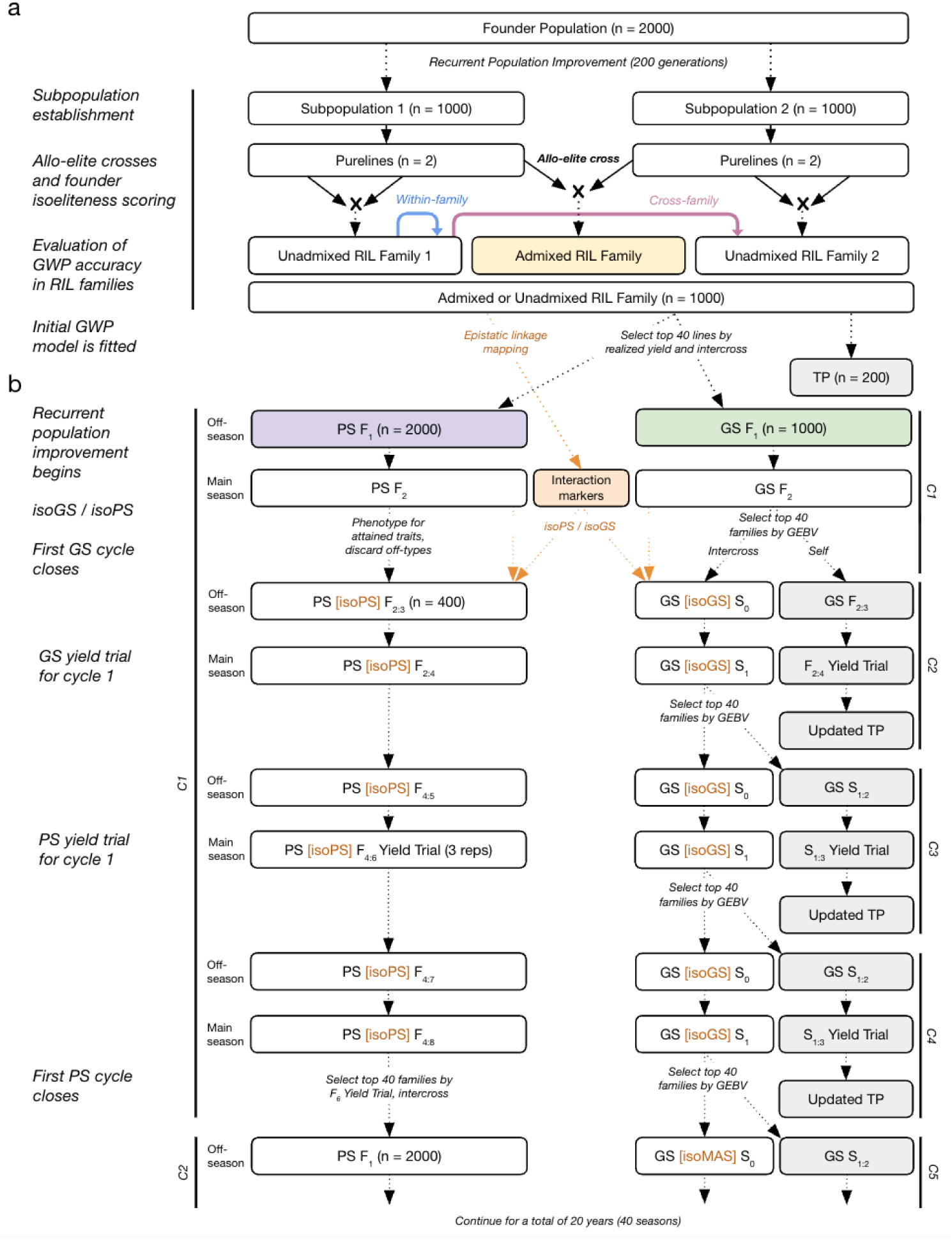
Framework of alternative recurrent population improvement breeding schemes. (a) Two independent subpopulations were derived from the same founder population, representative of the development of two breeding programs.Purelines from each subpopulation, assumed to be allo-elite, were crossed to create admixed RIL families (in addition to unadmixed RIL families derived from a cross between two lines from the same subpopulation). This enabled evaluation of 3 different GWP testing scenarios: “within-family”, “cross-family”, and “admixed”. (b) Each RIL family was then evaluated for genetic gain potential through four recurrent population improvement schemes for comparison: GS, PS, isoGS and isoPS. To develop each initial population, the top 40 RILs from each RIL family were intercrossed to create an F_1_ population, which was recurrently improved for 40 seasons, alternating between a “main season”, where phenotypes could be measured, and an “off season”, where they could not. isoGS and isoPS were applied to the F_2_ generation of cycle 1 only, after which the populations were improved through standard PS or GS.

### Calculating genetic variance of realized yield

Since GWP performance depends in part on genetic variance in the TP, the genetic variance of realized yield was calculated in each TP. The genetic value of realized yield for each individual was calculated based on the genetic values of *t_1_, t_2_, and t_3_*. The total genetic variance was calculated as the variance of the genetic values of realized yield. Correlations between mean isoeliteness, desired trait heterogeneity, log-transformed genetic variance, and *r*_MG_ were assessed with the pairs.panels() function from the R package “psych” (Revelle 2026). The partial correlation between *r*_MG_ and mean isoeliteness, controlling for log-transformed genetic variance and desired trait genetic heterogeneity, was calculated with the pcor.test() function from the R package “ppcor” (Kim 2015).

### Evaluating genetic gain through population improvement

To assess how genetic gain was impacted by the isoeliteness of the initial founders and different breeding strategies, several independent recurrent population improvement schemes were applied to each admixed and unadmixed RIL family (Fig. 2b). First, the top 40 RILs based on realized yield were selected for advancement. Since the mean realized yield of the 40 founders was correlated with long-term genetic gain, to enable a fair comparison between different levels of founder isoeliteness, the mean realized yield of the founders had to be controlled. A positive correlation was observed between the mean isoeliteness of the initial founders of the RIL families and the mean realized yield of the 40 top RILs. Simulation replicates were therefore filtered to retain only those with a mean realized yield across the top 40 RILs within 1 unit of the grand mean of realized yield before filtering (146). This eliminated the correlation between mean isoeliteness and mean initial realized yield, and left 691 replicates for analysis.

Recurrent population improvement was initiated by intercrossing the 40 selected RILs to generate an F_1_ population with 200 full-sibling families. Population improvement was carried out using both GS and PS for comparison. GS populations started with 5 seeds per cross (for a total of 1000 F_1_ plants), and PS populations started with 10 seeds per cross (for a total of 2000 F_1_ plants). Cycle length was 8 seasons (4 years) for PS and 2 seasons (1 year) for GS. Each year consisted of two seasons: an “off-season” (e.g. a winter nursery), where phenotypic data was not collected and lines were only selfed, and a “main-season” (e.g. a summer nursery), where phenotypic data could be collected and crosses could be made. Recurrent population improvement continued for a total of 40 seasons (i.e. 20 cycles of GS and 5 cycles of PS).

Genetic gain for each cycle was calculated as the difference in mean realized yield of the founders advanced to the next cycle compared to the mean realized yield of the founders of the current cycle. The Pearson correlation between founder isoeliteness and genetic gain after each cycle was calculated across all replicates, for each improvement scheme, with the cor() function from base R (R Core Team 2025). At the beginning of each cycle, the genetic variance among F_1_ (S_0_) lines for realized yield was calculated as described above.

### Phenotypic recurrent selection

Each PS cycle (Table 3) began in the off-season by selfing the F_1_ plants (Fig. 2b). Then, during the main season, the bottom 20% of the F_2_ population was discarded as off-types based on suitability, representative of the natural culling done by a breeder. For isoPS populations, the top 25% of F_2_ genotypes with ‘parent 1’ haplotypes were then selected, as described above. For a fair comparison, in PS populations, 25% of the F_2_ genotypes not discarded as off-types were randomly selected for advancement. The remaining 400 F_2_ genotypes were selfed. In the next off-season, the F_2:3_ genotypes were selfed, and in the subsequent main season, one random genotype from each of the 400 F_2:4_ plots was selected and selfed. These F_4:5_ lines were selfed in the next off-season, and in the following main season, F_4:6_ lines were phenotyped for realized yield in a three-replicate “yield trial”. In the following off-season, the F_4:7_ genotypes were again selfed, so that the crossing could happen in the subsequent main season. The top 40 F_4:8_ lines were selected based on the F_4:6_ yield trial, from which a random genotype per line was selected. 200 random crosses were made between these 40 lines to close the cycle.

**Table 3.** Phenotypic population improvement.

| **Season** | **Season Type** | **Stage** | **Plots** | **Plants per Plot** | **Families / Lines** | **Plants** | **Activity** |
|---|---|---|---|---|---|---|---|
| 1 | Off | F<sub>1</sub> | 200 | 10 | 200 | 2000 | Self all |
| 2 | Main | F<sub>2</sub> | 200 | 10 | 200 | 2000 | Select top 80%, then self top 25% of those |
| 3 | Off | F<sub>2:3</sub> | 400 | 20 | 400 | 8000 | Self all |
| 4 | Main | F<sub>2:4</sub> | 400 | 20 | 400 | 8000 | Self random plant per plot |
| 5 | Off | F<sub>4:5</sub> | 400 | 20 | 400 | 8000 | Self all |
| 6 | Main | F<sub>4:6</sub> | 1200<br>(3 reps per line) | 20 | 400 | 24000 | Yield trial |
| 7 | Off | F<sub>4:7</sub> | 400 | 20 | 400 | 800 | Self all |
| 8 | Main | F<sub>4:8</sub> | 400 | 20 | 400 | 8000 | Make 200 crosses |

### GS-based recurrent selection

For the GS pipeline (Table 4), the initial TP for the RR-BLUP model was the 40 RIL founders, plus 160 randomly selected RILs meant to capture the genetic diversity of the RIL family, for a total of 200 lines. Each GS cycle began in the same way as the PS cycles (Fig. 2b), by selfing the F_1_ plants in the off-season, and discarding the bottom 20% of the F_2_ lines based on suitability. The isoGS method was then used to select the top 25% of genotypes. From the remaining genotypes, the top 40 F_2_ families were selected in the main season based on GEBV’s produced by the initial RR-BLUP model. The top genotype by GEBV from each family was advanced, and 200 random crosses between the 40 lines were made to produce 200 S_0_ families and close the cycle. GWP accuracy (*r*_MG_) was calculated for the 40 selected lines at each cycle. Additionally, one sibling of each of the selected 40 lines was randomly selected, and selfed and bulked for two more seasons to produce 40 F_2:4_ lines, which were phenotyped for realized yield. The GWP TP was updated using a random genotype from each of the 40 F_2:4_ lines, plus the top 160 lines from the existing TP based on realized yield. In cycle 2, the target population was therefore two cycles removed from the TP, while in cycle 3 and subsequent cycles, the target population was only one cycle removed from the TP. As negative controls, GS and isoGS were also carried out without updating the initial TP.

**Table 4.** GS-based population improvement.

| Season | Season Type | Stage | Plots | Plants per Plot | Families / Lines | Plants | Activity |
|---|---|---|---|---|---|---|---|
| 1 | Off | F<sub>1</sub> (S<sub>0</sub>) | 200 | 10 | 200 | 2000 | Self all |
| 2 | Main | F<sub>2</sub> (S<sub>1</sub>) | 200 | 10 | 200 | 2000 | Select top 80%, then self top 25% of those |
| 3 | Off | F<sub>3</sub> (S<sub>1:3</sub>) | 40 | 20 | 400 | 8000 | Self all |
| 4 | Main | F<sub>4</sub> (S<sub>1:4</sub>) | 120<br>(3 reps per line) | 20 | 400 | 8000 | Self random plant per plot |

### Assessing cost-effectiveness of each breeding scheme

To enable a practical comparison between schemes for return on investment, each scheme was assessed according to its “cost-per-unit-gain”. Costs of RIL family development and epistatic linkage mapping were not included, as analysis was focused on activities related directly to population improvement. Genotyping costs were based on current estimates from the CGIAR Excellence in breeding toolbox: (https://excellenceinbreeding.org/toolbox/collections/5). Mid-density Diversity Array Technology genotyping (DArTag) at $10/sample was used for GWP and Kompetitive Allele-specific PCR (KASP) marker genotyping at $5/sample was used for isoGS and isoPS. Additionally, a “low sequencing cost” was evaluated, assuming a future reduction in genotyping costs of 80% (with DArTag and KASP costs of $2/sample and $1/sample, respectively).

For labor costs, including population establishment, field maintenance, harvesting, tissue collection, selfing and crossing, two scenarios were considered. A “low labor cost” scenario was based on cost estimates from the Savannah Agricultural Research Institute in Ghana, and a “high labor cost” scenario was based on estimates from Kansas State University (Table 3). For a full breakdown of costs, see Supplementary Tables 1 and 2.

## RESULTS

### Isoeliteness was predictive of admixed GWP accuracy

To confirm the difficulty of using GS in heterogeneous germplasm in our simulations, the accuracy of GWP was evaluated in three target population scenarios defined by their relationship to the TP (Fig. 2a). As expected, the “within-family” scenario, representative of standard use of GWP in a closed breeding population, had the highest accuracy (mean *r*_MG_ = 0.62) (Fig. 3a). The “cross-population” scenario, included as a negative control, had a mean *r*_MG_ of 0.034, consistent with expectations of reduced GWP accuracy when the target population has little kinship with the TP (Habier et al. 2007). The “admixed” scenario, representative of an open or genetically heterogeneous breeding population, had a mean *r*_MG_ of 0.57, though there was considerable variability. Considering this admixed scenario only, there was a strong positive correlation (*r* = 0.77, *p*-value < 0.001) between mean isoeliteness of the RIL founder lines and GWP accuracy (Fig. 3b). Additionally, there was also a strong negative correlation (*r* = −0.66, *p*-value < 0.001) between mean *r*_MG_ and the log genetic variance of breeding fitness, as well as a weak correlation (*r* = −0.08, *p*-value < 0.001) between mean *r*_MG_ desired trait (e.g. yield potential) genetic heterogeneity (Supplementary Fig. 1). When these two factors were controlled for with a partial correlation test, there was still a strong correlation of *r* = 0.51 (*p*-value < 0.001) between mean isoeliteness and mean *r*_MG_. There was no correlation between mean isoeliteness and desired trait heterogeneity (Supplementary Fig. 2). This validated that the low predictive accuracy of the GWP models in diverse germplasm was primarily due to newly segregating attained trait QTL in the RIL families.

**Figure 3.**
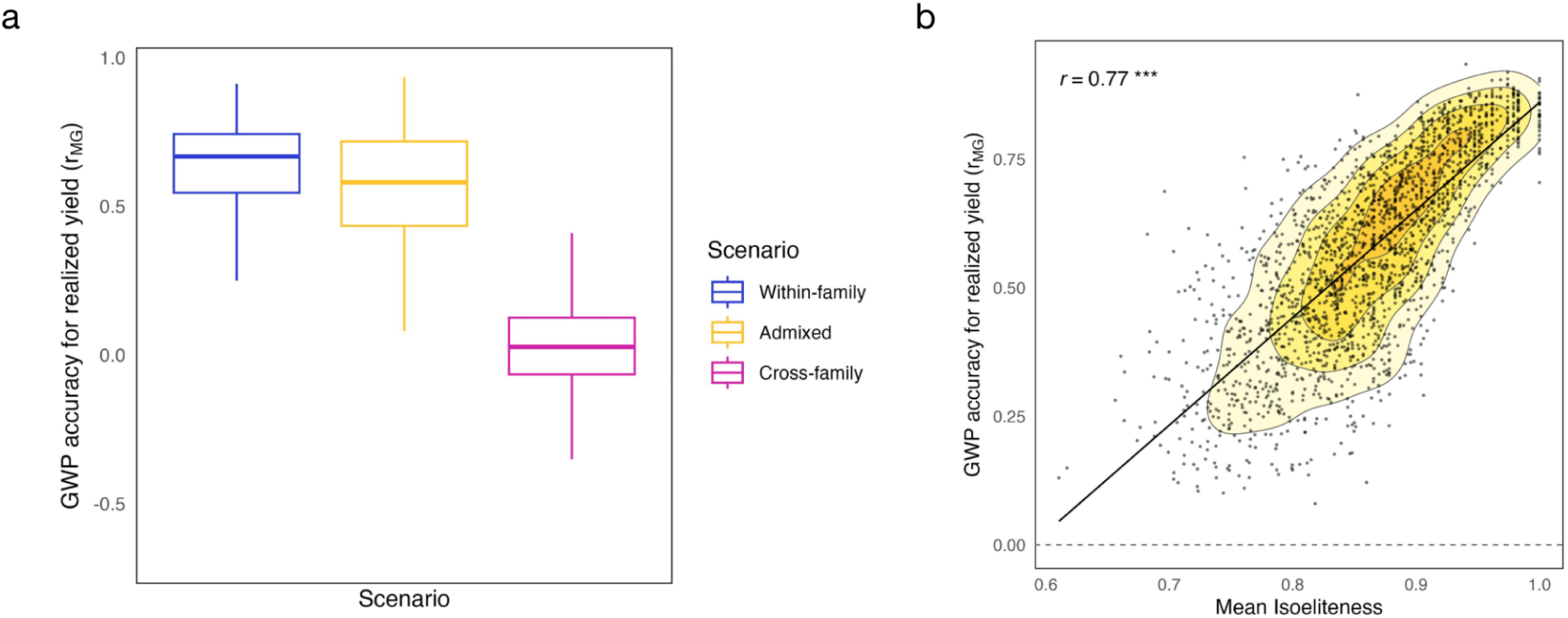
GWP accuracy was higher in admixed families derived from parents with high isoeliteness. An RR-BLUP model was trained in five different scenarios: within-family, admixed, and cross-family (as a negative control). (a) GWP accuracy (*r*_MG_) in each scenario matched expectations, as within-family was the highest, admixed was slightly lower, and cross-family was the lowest. (b) For the admixed scenario, there was a positive correlation (*r*) of 0.77 between mean isoeliteness of the parents and GWP accuracy.

### Impact of founder isoeliteness on genetic gain

Next, the implications of founder isoeliteness on genetic gain were evaluated under different GS and PS schemes, using populations derived from intercrossing the top 40 RILs in each RIL family. Recurrent population improvement was simulated for 40 seasons (i.e. non-overlapping generations), corresponding to 20 years, 20 cycles of GS and 5 cycles of PS. Each population was categorized based on the isoeliteness of the parents of the RIL families from which they were derived as low (≤ 0.8), moderate (between 0.8 and 0.9), or high (≥ 0.95). Among populations with high founder isoeliteness, there was substantially higher genetic gain in populations derived from admixed families than in populations derived from unadmixed families, reflecting the wider availability of genetic variation. This was true both for GS (151%, 269% and 251% higher after 4, 12, and 20 cycles), and for PS (108%, 130% and 150% higher after 4, 12, and 20 cycles) (Fig. 4). Subsequent analysis was only done on populations derived from admixed families. In populations with high founder isoeliteness, GS led to higher genetic gain than PS (58% higher after 4 cycles and 23% higher after 12 cycles), though PS surpassed GS by cycle 20 (Fig. 4).

**Figure 4.**
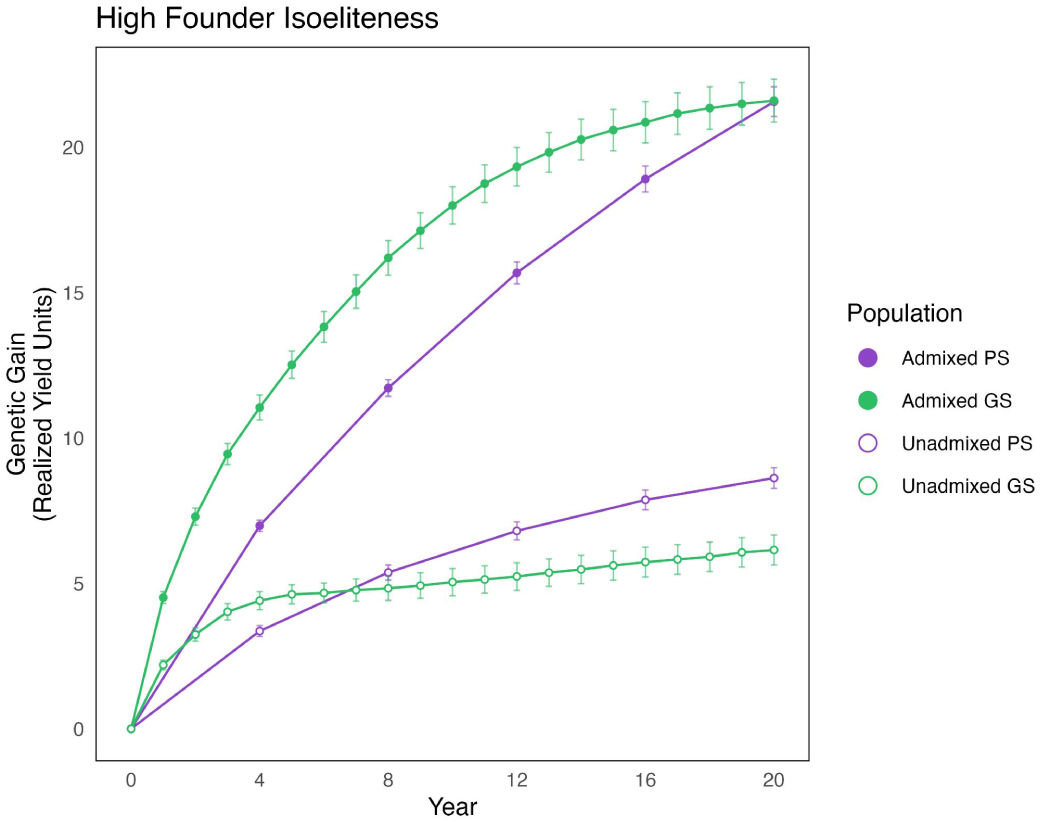
Admixed populations with high founder isoeliteness had more genetic gain than unadmixed populations. Considering only populations with high founder isoeliteness, the mean genetic gain at each cycle was aggregated by selection scheme, and plotted along with 95% confidence interval bars. There was more genetic gain in admixed populations than in unadmixed populations, and higher initial genetic gain from GS than from PS. In the long-term, PS surpassed GS in both admixed and unadmixed populations.

The effect of founder isoeliteness on genetic gain differed between GS and PS schemes, primarily in early cycles. Genetic gain with GS was 63% lower after 2 cycles and 27% lower after 4 cycles in populations with low founder isoeliteness than in populations with high founder isoeliteness (Fig. 5a, Supplementary Table 3). Notably, there was a decrease in genetic gain for populations with low founder isoeliteness in cycle 2, which were two cycles removed from the initial TP. This reduced genetic gain was likely due to the low GWP accuracy in early cycles of populations with low founder isoeliteness (mean *r*_MG_ = 0.13 in cycle 2 compared to mean *r*_MG_ = 0.28 and 0.49 in populations with moderate and high founder isoeliteness, respectively) (Supplementary Fig. 2). Populations with moderate founder isoeliteness had similar genetic gain as populations with high founder isoeliteness until cycle 12, after which they had higher gain. Populations with low founder isoeliteness also caught up to populations with moderate founder isoeliteness by cycle 12. The differences in genetic gain between different levels of founder isoeliteness were most striking when the initial GWP model was not updated after being fitted to the RIL TP (Supplementary Fig. 3). By cycle 20, populations with low founder isoeliteness without an updated GWP model had a 85% reduction in genetic gain, while populations with high founder isoeliteness had a 32% reduction in genetic gain.

**Figure 5.**
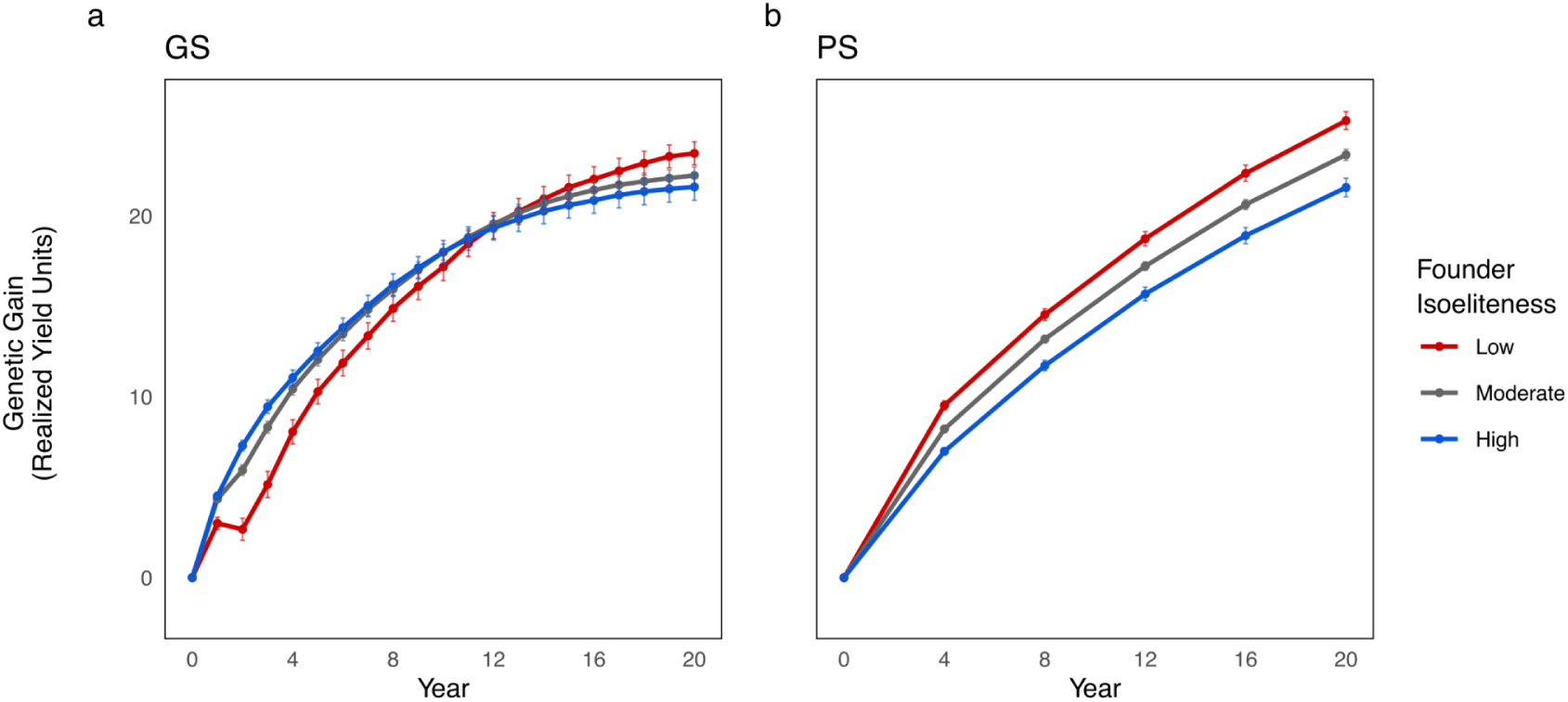
GS was ineffective in early cycles for populations with low founder isoeliteness. Genetic gain was aggregated by population improvement scheme and founder isoeliteness category. The mean genetic gain in “realized yield” units, along with 95% confidence intervals across 691 replicates is shown. (a) With GS, populations with high founder isoeliteness had smooth and rapid genetic gain, while populations with low and moderate founder isoeliteness had slower genetic gain in early cycles. Genetic gain was negative in year 2 for populations with low founder isoeliteness. By year 20, genetic gain was highest in populations with low founder isoeliteness, followed by populations with moderate and high founder isoeliteness populations. (b) With PS, the highest genetic gain occurred in populations with low founder isoeliteness, followed by populations with moderate and high founder eliteness.

In striking contrast with its effect in GS schemes, low founder isoeliteness had a positive effect on genetic gain in early cycles in PS schemes (Fig. 5b). Populations with low founder isoeliteness had 36% higher genetic gain after 4 years (1 cycle) than populations with high founder isoeliteness, and maintained higher genetic gain throughout all 20 years. By year 20, in both GS and PS, populations with low and moderate founder isoeliteness had higher genetic gain than populations with high founder isoeliteness, highlighting the potential long-term benefit of crossing parents with low isoeliteness.

### isoGS led to increased genetic gain in populations with low and moderate founder isoeliteness

To mitigate the potential detrimental effects of fitness epistasis among the attained trait QTL, isoGS and isoPS were used to recover F_2_ genotypes with parental haplotypes at loci that colocalized with significant detected fitness interactions (Fig. 1). Populations were then improved through standard PS or GS for the remaining cycles. isoGS provided the most benefit in low and moderate founder isoeliteness populations, particularly in early cycles (Supplementary Fig. 4), and improved long-term genetic gain in most scenarios (Fig. 6). In populations with low founder isoeliteness, isoGS led to 128% higher genetic gain than naive GS after 2 years, 31% higher gain after 4 years, and 15% higher gain after 20 years (Fig. 6a, Supplementary Table 3). The improvement in GWP accuracy in isoGS compared to GS further validated the effectiveness of the strategy. In populations with low founder isoeliteness, mean *r*_MG_ was 103% higher in cycle 2 and 25% higher in cycle 4 (Supplementary Fig. 2). isoPS had a smaller relative benefit, as naive PS already benefitted from low founder isoeliteness (Fig. 5b). In populations with low founder isoeliteness, genetic gain with isoPS was just 8% higher compared to naive PS after 4 years and identical after 20 years (Fig 6a, Supplementary Table 3). In populations with high founder isoeliteness populations, isoGS and isoPS did not provide any benefit, and GS strongly outperformed PS until later cycles (Fig. 6c).

**Figure 6.**
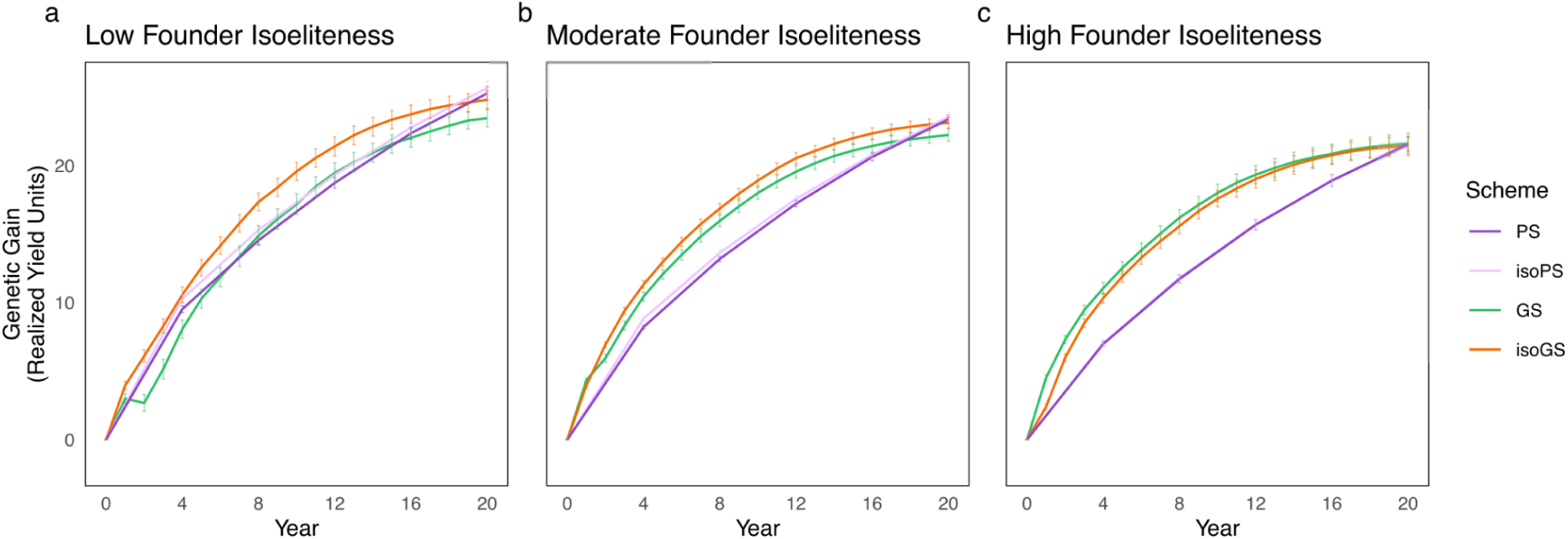
isoGS was most effective in populations with low founder isoeliteness. The 691 replicates were aggregated by founder isoeliteness category, to compare the genetic gain achieved through each population improvement scheme. (a) For populations with low founder isoeliteness, isoGS had the highest genetic gain until year 18, and isoPS outperformed PS. (b) For populations with moderate founder isoeliteness, isoGS and GS led to more genetic gain than PS or isoPS in early cycles, and isoGS was higher than GS throughout all cycles. (c) For populations with high founder isoeliteness, GS and PS outperformed isoGS and isoPS, respectively. Naive GS led to the most genetic gain, and had a larger advantage over PS compared to populations with low and moderate founder isoeliteness.

Across all improvement methods, the correlation between founder isoeliteness and genetic gain decreased with each cycle (Supplementary Fig. 5), indicating the diminishing importance of founder isoeliteness in later cycles. Consistent with the genetic gain findings, the correlation was highest with naive GS, decreased fastest with isoGS, and was consistently negative for PS and isoPS. There were insignificant differences found when isoGS was used with perfect markers, though the low-density markers were less effective than the high density markers at driving early genetic gain (Supplementary Fig. 3). Additionally, founder isoeliteness impacted the relative cost-effectiveness of each improvement scheme in terms of cost-per-unit gain. For populations with low founder isoeliteness, isoGS was more cost-effective than GS in early cycles, while in high founder isoeliteness populations, naive GS was more cost-effective (Fig. 7a). GS was more cost effective than PS only under assumptions of high labor costs (Fig. 7b) or low sequencing costs (Supplementary Fig. 6). Since PS was insensitive to founder isoeliteness, isoPS was an unnecessary added cost in all scenarios. Taken together, these findings demonstrate the value of quantifying founder isoeliteness for informing population improvement strategies, especially in programs that utilize GS.

**Figure 7.**
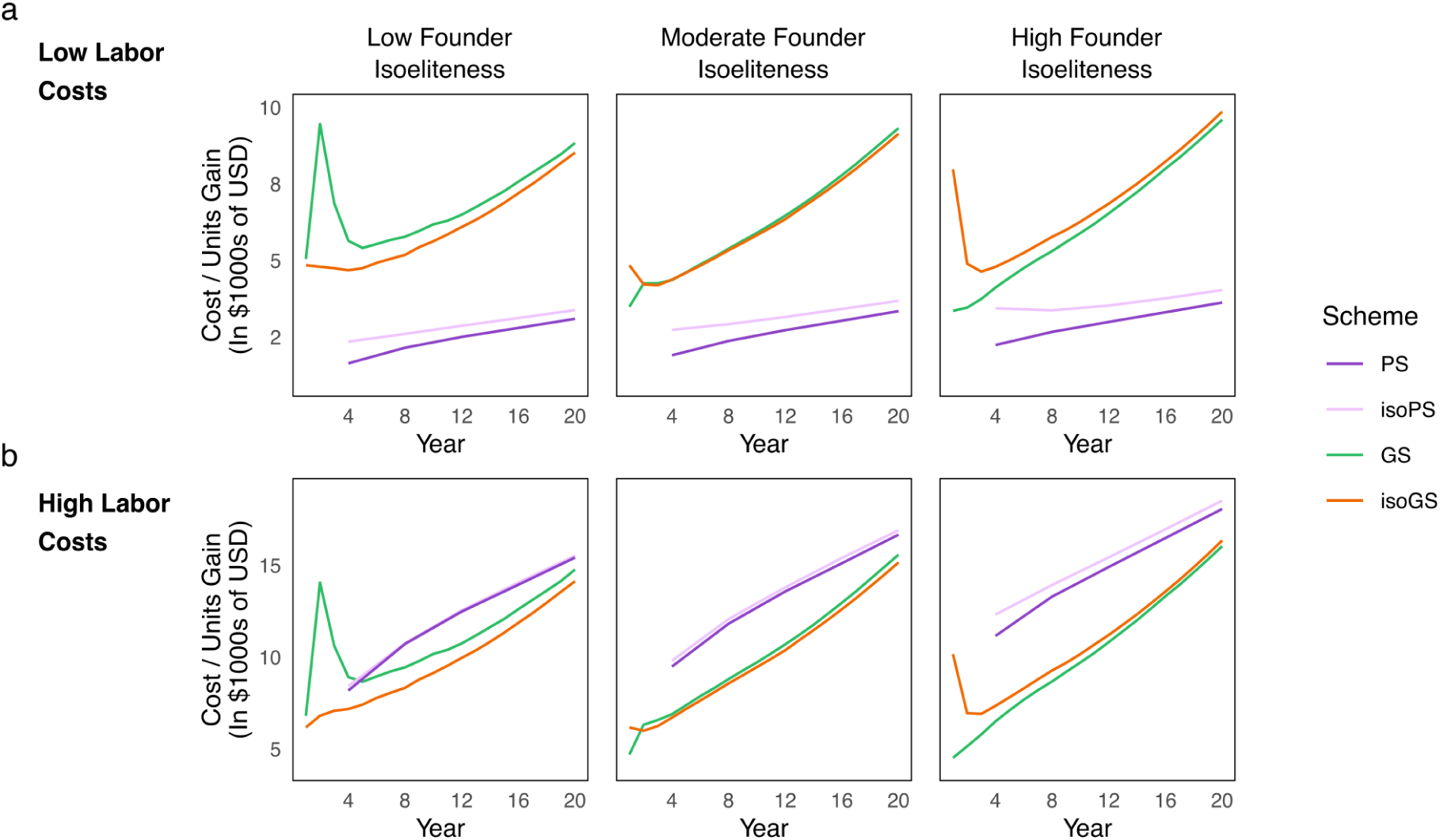
The cost effectiveness of each scheme was dependent on founder isoeliteness and labor costs. Cumulative costs per unit of genetic gain (in $1000s of USD) are shown for GS, isoGS, PS, and isoPS, under two contrasting labor cost assumption scenarios. (a) Under assumptions of low labor costs, PS and isoPS were much more cost-effective than GS or isoGS. isoGS was more cost-effective than GS in early cycles in populations with low founder isoeliteness, and became relatively cost-ineffective in populations with high founder isoeliteness. The cost-effectiveness of GS was closest to that of PS in populations with high founder isoeliteness. (b) Under high labor cost assumptions, isoGS was the most cost-effective scheme for populations with low founder isoeliteness, while GS was most effective in populations with high founder isoeliteness. GS was generally more cost-effective than PS across all levels of founder isoeliteness.

## DISCUSSION

### Crossing genetically diverse lines creates higher genetic gain potential

The premise of the breeder’s dilemma between diversity and eliteness is that there is inherent value in having genetic variation to drive continued genetic gain for desired traits (Falk 2010; Cobb, Juma, et al. 2019; Yang et al. 2020; Anderson et al. 2021; Swarup et al. 2021). The substantial increase in genetic gain in admixed populations relative to unadmixed populations (Fig. 4) validated this upside. Furthermore, the higher genetic gain obtained in populations with low founder isoeliteness improved through isoGS, PS, and isoPS (Fig. 5), and the low correlation between founder isoeliteness and desired trait genetic heterogeneity (Supplementary Fig. 1), suggests that a release of genetic variation underlying attained traits could even lead to improved attained trait phenotypes compared to the founders (Zebell et al. 2025). These findings are consistent with the motivations of the intentional introduction of novel genetic variation into a closed breeding population through pre-breeding (Dudley 1982; Simmonds 1993; Hallauer and Carena 2012).

Our assumption of two essential attained traits with high heritability, with equivalent and symmetrical penalties on realized yield was simplified compared to an actual breeding scenario. In reality, an ideotype may consist of many attained traits, some of which may be difficult to phenotype in early generations, in which case maintaining eliteness in an open breeding population would have higher complexity than in our simulations (Rasmusson and Phillips 1997). It also is worth noting that our PS scheme, in which F_6_ lines were evaluated in year 3 yield trials, and cycles were closed in year 4, was highly aggressive and perhaps unrealistic (Hickey et al. 2017; Cobb, Juma, et al. 2019; Rutkoski 2019a). This short cycle length was chosen to enable our PS scheme to be competitive with our similarly aggressive 1-year cycle GS scheme. It would be worthwhile to validate our findings using empirical data or a different set of parameters.

### isoGS effectively converts fitness epistasis to selectable additive variation

In spite of the benefits of genetic diversity for genetic gain, it is typically advantageous to maintain closed breeding populations, in part to retain favorable co-adapted epistatic complexes (Rasmusson and Phillips 1997; Condón et al. 2009; Falk 2010; Tessele et al. 2026). However, nascent breeding programs, still developing their elite gene pools, may have genetically diverse founder lines. Additionally, breeding populations are often re-opened to introgress abiotic tolerance and biotic stress resistance traits from other elite gene pools (Bailey-Serres et al. 2010; Boukar et al. 2020; Dormatey et al. 2020; Hernandez et al. 2020), which may be allo-elite (Fall et al. 2026). In these circumstances, fitness epistasis is expected to be pervasive when allo-elite crosses disrupt co-adapted epistatic complexes (Wright 1935; Whitlock et al. 1995; Brodie 2000; Monyak and Morris 2026). Our results suggest that GS can be highly ineffective in early cycles of populations derived from founders with low isoeliteness (Fig. 5a), while PS is less impacted (Fig. 5b). Furthermore, while our simulations considered purely additive traits where epistasis only emerges at the fitness level, biological epistasis is expected to be present in diverse populations (Podlich and Cooper 1999; Holland 2001; Carlborg et al. 2006), suggesting our results may underestimate the importance of epistasis for breeding network operations.

Standard GWP models such as RR-BLUP (Meuwissen et al. 2001) and GBLUP (Bernardo 1994) are designed for use with traits controlled by numerous small-effect additive QTL (Crossa et al. 2017). While methods such as LASSO (Tibshirani 1996) and Bayes-B (Wimmer et al. 2013) can better accommodate large-effect variants, such as in the oligogenic traits we modeled, these methods are not expected to be able to handle epistasis. The strong association between isoeliteness and *r*_MG_ in the admixed RIL families (Fig. 3b, Supplementary Fig. 1) suggests that segregating epistatic haplotypes made it difficult for the initial GWP model to estimate accurate marker effects. Since there is also expected to be weak linkage disequilibrium between markers and causal loci in an admixed family, the population structure was likely a factor as well (Habier et al. 2010; Jannink 2010; Windhausen et al. 2012; Bernardo 2020; Bernardo 2025). The low *r*_MG_ (Supplementary Figs. 2a-b) and genetic gain (Fig. 5a) in early improvement cycles of populations with low and moderate founder isoeliteness further validated the epistasis hypothesis, since major-effect beneficial alleles in the RIL TP acting in an additive manner should have been enriched through naive GS. The fact that the TP was not updated until year 2 (Fig. 2b) likely exacerbated this phenomenon. The use of isoGS led to a striking improvement in *r*_MG_ in early cycles (Supplementary Figs. 2a-b) and a corresponding improvement in genetic gain (Figs. 6a-b, Supplementary Fig. 4). By facilitating the fixation of favorable attained trait haplotypes in the first generation of selection, isoGS converted epistatic variation into selectable additive variation (Carlborg et al. 2006; Technow et al. 2021; Ortiz-Barrientos et al. 2026), enabling faster genetic gain.

### Practical considerations for breeding programs considering isoGS

GS offers numerous potential advantages over PS (Meuwissen et al. 2001; Bernardo and Yu 2007; Heffner et al. 2010; Hickey et al. 2017), but our findings implicate founder isoeliteness as a key factor in determining whether these benefits can be realized. Our results suggest that when founders with high isoeliteness can be identified, the resulting populations can be directly pushed for genetic gain through rapid cycling with GS without consideration for fitness epistasis in the population (Figs. 4, 6c). In contrast, PS was insensitive to different founder isoeliteness levels, and as founder isoeliteness increased, the benefit of GS over PS decreased accordingly (Fig. 6). These insights may also be relevant for the prospects of GS-enabled pre-breeding (Bernardo 2009; Gorjanc et al. 2016; Allier et al. 2020; Sanchez et al. 2023). The strong performance of PS in populations with low founder isoeliteness (Fig. 5b), and the absence of an added benefit of isoPS (Fig. 6a) suggest that the top inbreds at the end of each cycle PS may have recovered favorable attained trait haplotypes as a byproduct of selection. Additionally, the plateauing performance of GS in the long-term could be due to the fact that our simulation framework did not leverage strategies such as weighted genomic selection (Goddard 2009), optimal haplotype stacking (Kemper et al. 2012), or optimal haploid value selection (Daetwyler et al. 2015) to prevent GS from imposing its characteristic genetic bottleneck and exhausting available genetic variation (Jannink 2010). Indeed, the genetic variance decreased more rapidly with GS than with PS (Supplementary Fig. 7), which likely explains why PS was the superior scheme in the long-term.

From a cost-effectiveness standpoint, the decision of whether to use isoGS or isoPS depends on a program’s operational costs and particular breeding scheme (Heffner et al. 2010; Krchov and Bernardo 2015; Muleta et al. 2019; Gholami et al. 2021). The purpose of this study was not to directly compare the costs of GS and PS to provide universal recommendations, but to provide a general framework for understanding, quantifying, and managing isoeliteness. In programs with low labor costs, it is difficult to justify the use of GS in most contexts under existing sequencing costs (Fig. 7a), especially when continuous retraining of GWP models is required (Bernardo 2025). However, under assumptions of higher labor costs (Fig. 7b), and future scenarios where the cost of sequencing continues to decrease (Supplementary Fig. 6), the cost-benefit analysis shifts more in favor of isoGS and GS. Additionally, for crops and programs where doubled haploids (Forster et al. 2007; Gaynor et al. 2017) or speed breeding (Watson et al. 2019) are available, GS can become an even more attractive option still. In such cases, when epistatic linkage mapping reveals significant interaction effects between alleles segregating in a genetically heterogeneous population, our simulations suggest that usage of isoGS is beneficial. Whether through selection of genetically compatible crosses or progeny, isoeliteness is a key determinant of breeding outcomes in genetically diverse populations.

## DATA AVAILABILITY

All code and data, including scripts to reproduce the simulations are published to Github at https://github.com/CropAdaptationLab/FitnessLandscapes.

## SUPPLEMENTARY FIGURES

**Supplementary Figure 1.**
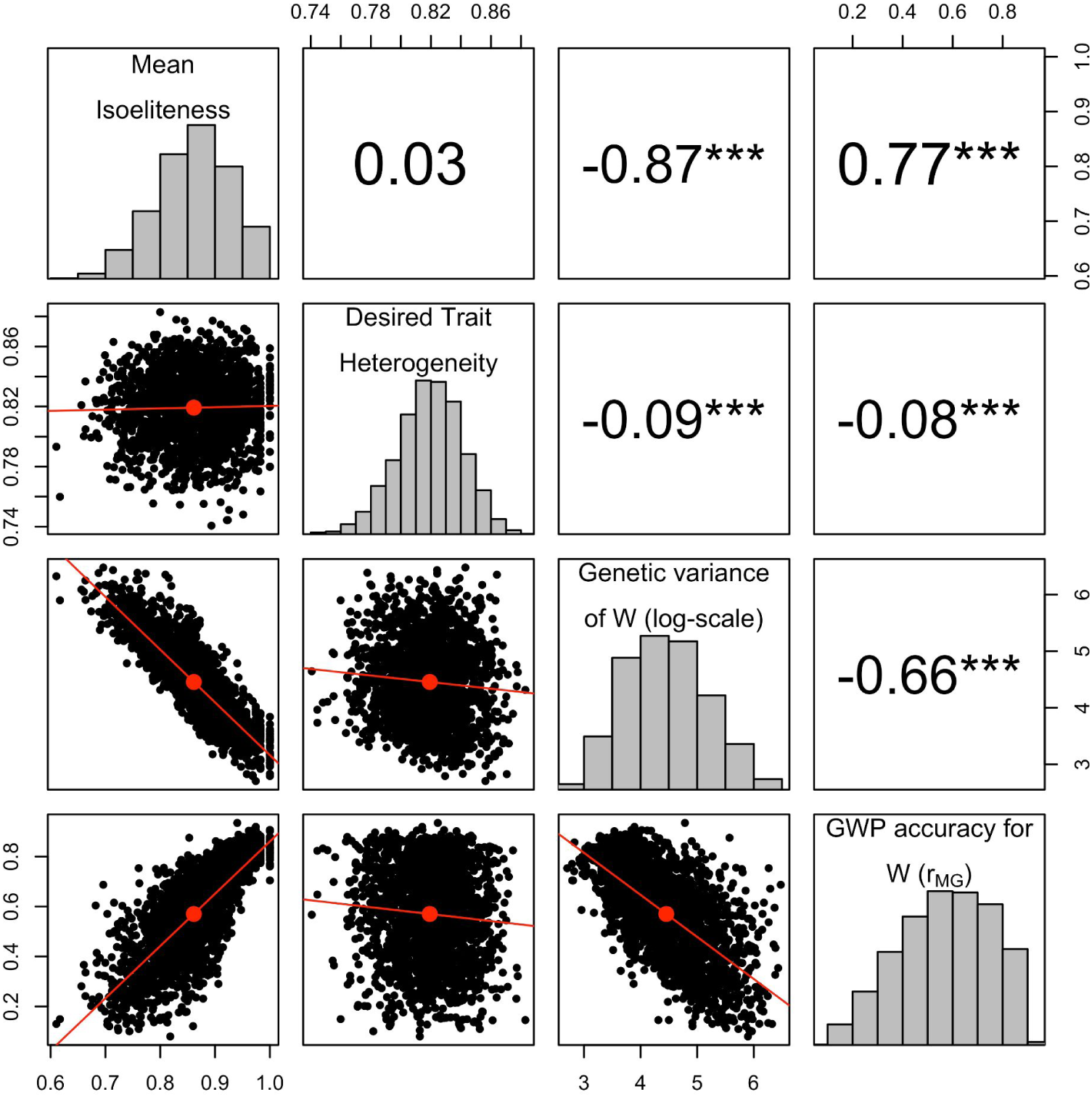
Factors associated with GWP accuracy in RIL families. For the admixed populations, “desired trait heterogeneity” and log genetic variance of *W* (realized yield) were calculated in addition to mean isoeliteness. Pearson correlations were calculated between these two metrics, as well as mean isoeliteness and GWP accuracy. There was no correlation between desired trait genetic heterogeneity and mean isoeliteness, and a weak correlation between desired trait genetic heterogeneity and GWP accuracy. There was a strong correlation between log genetic variance and mean isoeliteness, and between log genetic variance and GWP accuracy.

**Supplementary Figure 2.**
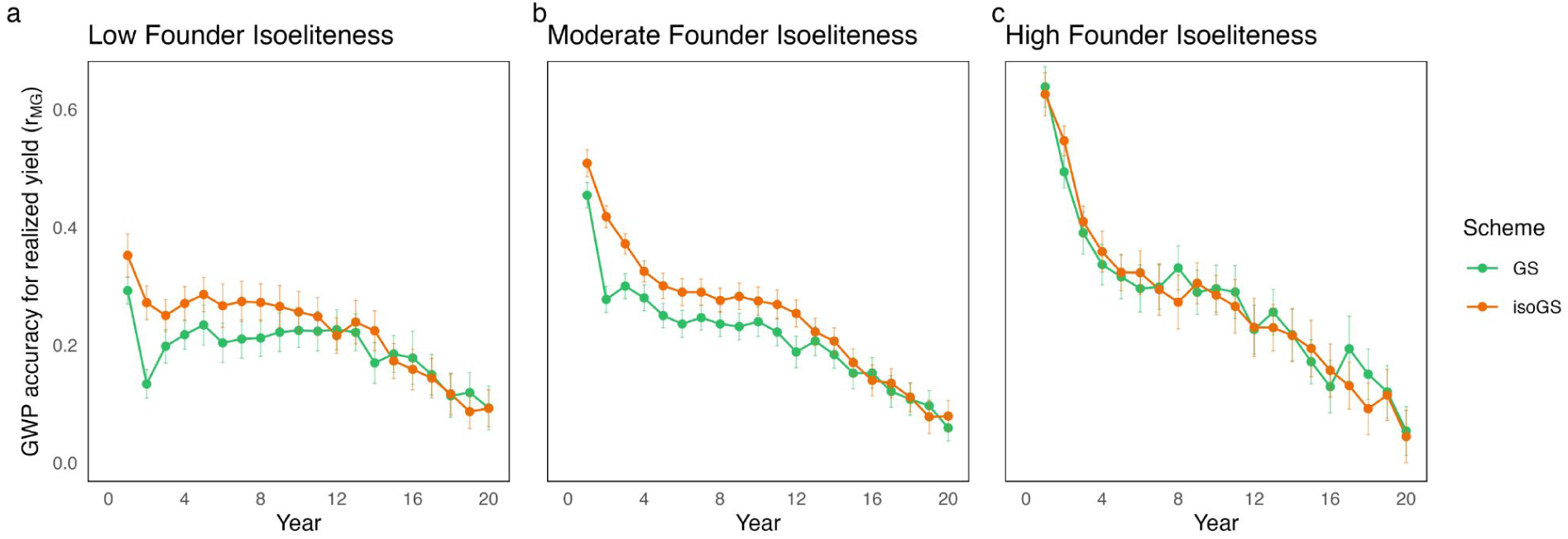
GWP accuracy in early cycles was affected by founder isoeliteness. The mean GWP accuracy (*r*_MG_) per year (with 95% confidence intervals) for GS and isoGS, grouped by founder isoeliteness, is shown. (a) For populations with low founder isoeliteness populations, accuracy with GS was low in early cycles, consistent with the negative genetic gain in early cycles (Fig. 5a). Accuracy was higher in isoGS populations. (b) Populations with moderate founder isoeliteness had higher initial GWP accuracy for GS in early cycles than populations with low founder isoeliteness, and isoGS had higher accuracy still. (c) In populations with high founder isoeliteness, GS and isoGS had near-identical GWP accuracy, which was higher in early cycles than in populations with low and moderate founder isoeliteness. Across all scenarios, GWP accuracy generally decreased with each cycle.

**Supplementary Figure 3.**
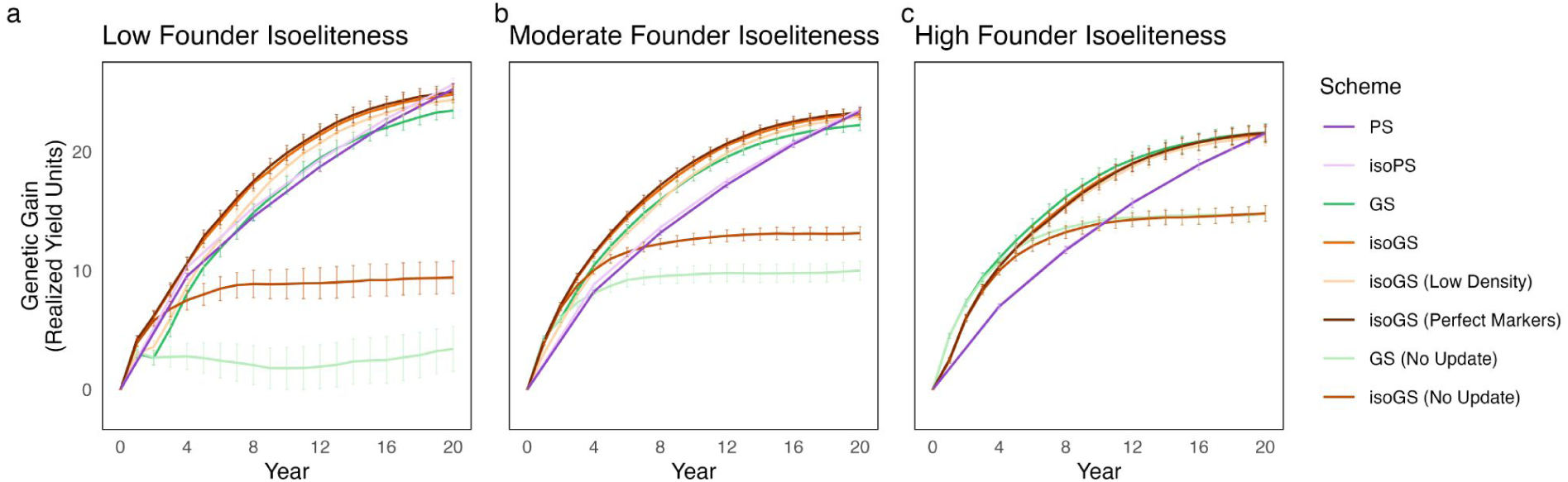
Positive and negative controls validate results. As negative controls, isoGS and GS populations were developed without updating the initial GWP model. This resulted in stagnant genetic gain in GS populations with low founder eliteness, which was moderately improved with isoGS (a). Populations with moderate and high founder isoeliteness (b, c) were less sensitive to not updating the model. isoGS was also performed with perfect markers (corresponding to the actual QTL) and lower density markers. There was little difference between the high-density markers used in the main simulations and the perfect markers, and a slight reduction in gain when using the low-density markers.

**Supplementary Figure 4.**
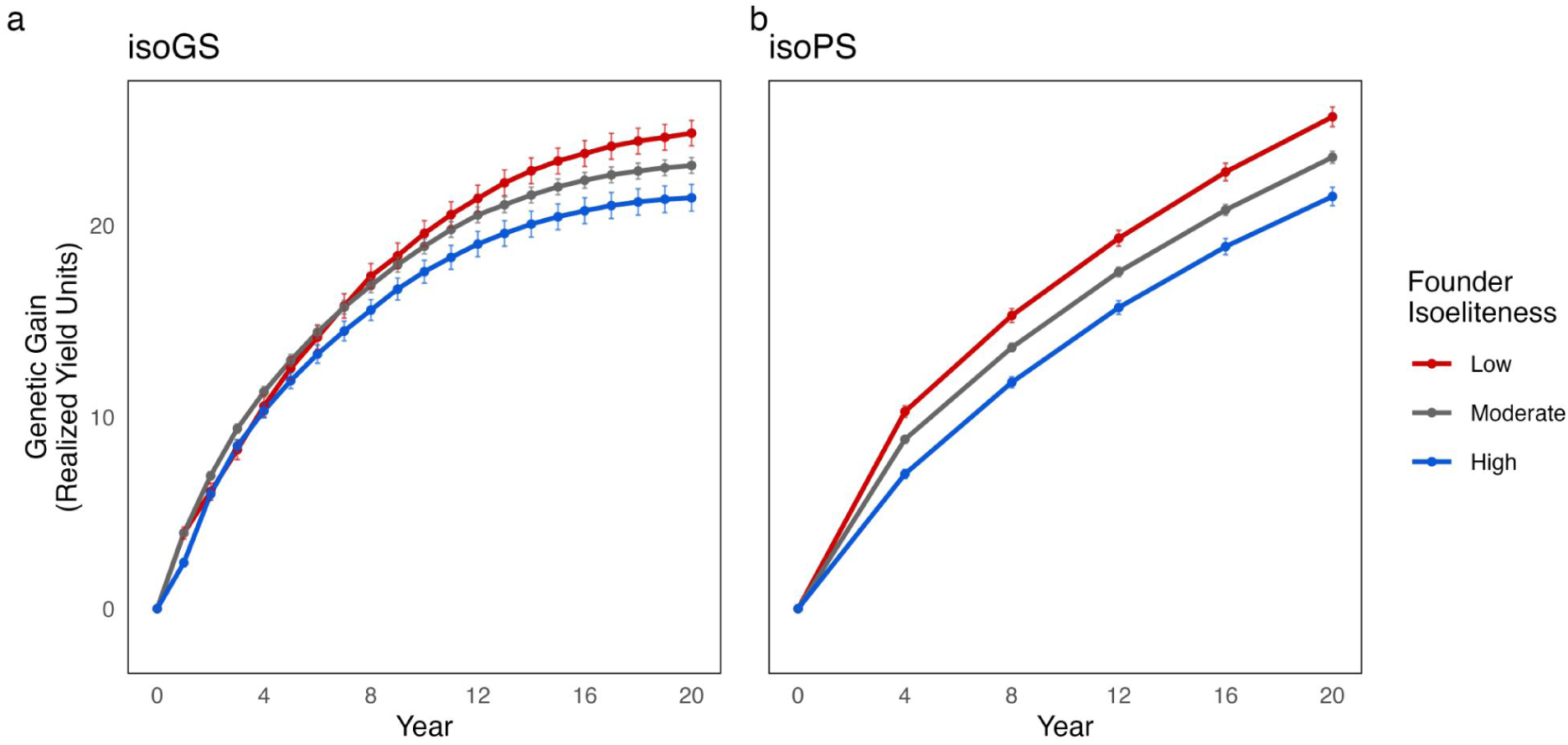
isoGS led to higher genetic gain in populations with low and moderate founder isoeliteness. With isoGS, the reduction in genetic gain in populations with low and moderate founder isoeliteness improved with GS (Fig. 5a) was eliminated (a). These populations had smooth genetic gain in early cycles and higher genetic gain than populations with high founder isoeliteness. (b) Though PS populations were already insensitive to low and moderate founder isoeliteness, isoPS led to even more genetic gain for these two scenarios.

**Supplementary Figure 5.**
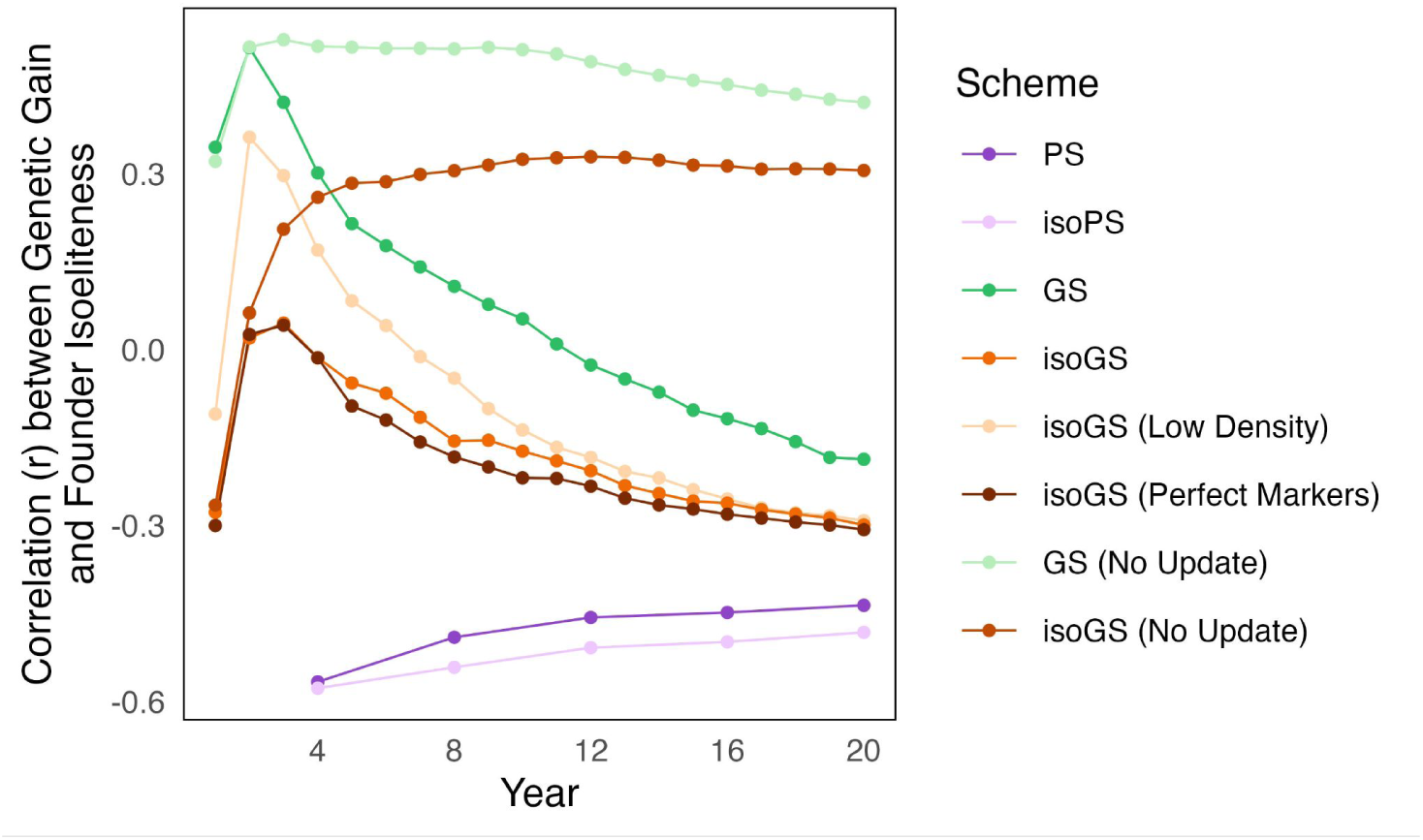
Correlation between founder isoeliteness and realized yield over time. The Pearson correlation (*r*) between founder isoeliteness and genetic gain was plotted by selection scheme, by year. The correlation for all GS populations decreased, indicating a decreased significance of founder isoeliteness over time. isoGS led to a lower correlation than naive GS, validating its effectiveness at negating the effects of low founder isoeliteness. isoPS and PS populations both had consistent negative correlations, consistent with the observation that isoPS and PS delivered more genetic gain in the populations with low founder isoeliteness. The two schemes that did not update the GWP model throughout had a consistently high correlation, which corresponded to their poor performances in populations with low and moderate founder isoeliteness.

**Supplementary Figure 6.**
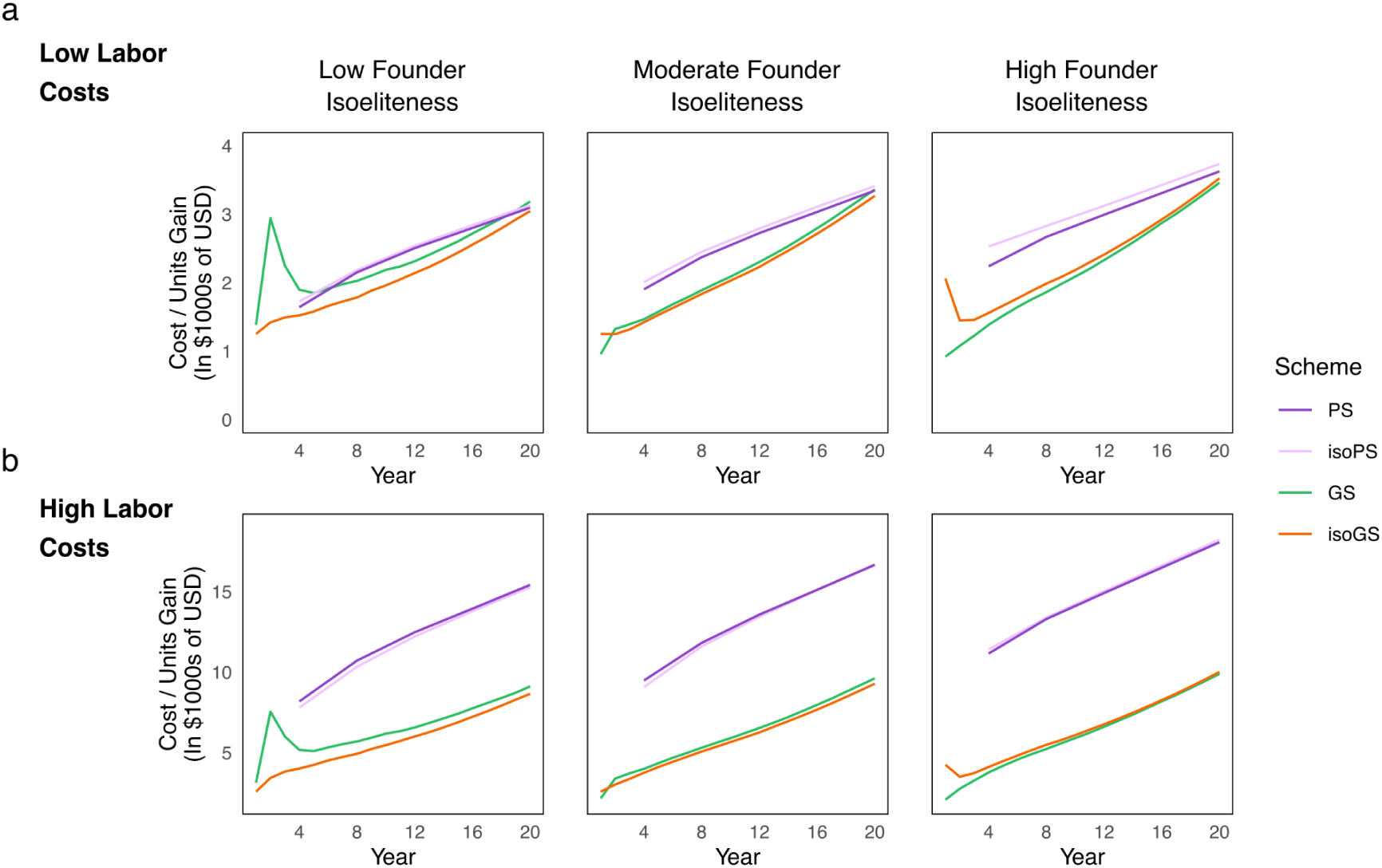
Cost analysis across future assumptions of sequencing costs. Cumulative costs per unit of genetic gain (in $1000s of USD) are shown for GS, isoGS, PS, and isoPS, under the assumption that the future cost of sequencing will decline by 80%. Two contrasting labor assumptions are shown, as in Fig. 7. GS was more effective than PS across all levels of founder isoeliteness and assumptions about labor. isoGS was the most cost-effective scheme in populations with low founder isoeliteness under both labor cost assumptions. For populations with high founder isoeliteness, GS was the most cost-effective scheme.

**Supplementary Figure 7.**
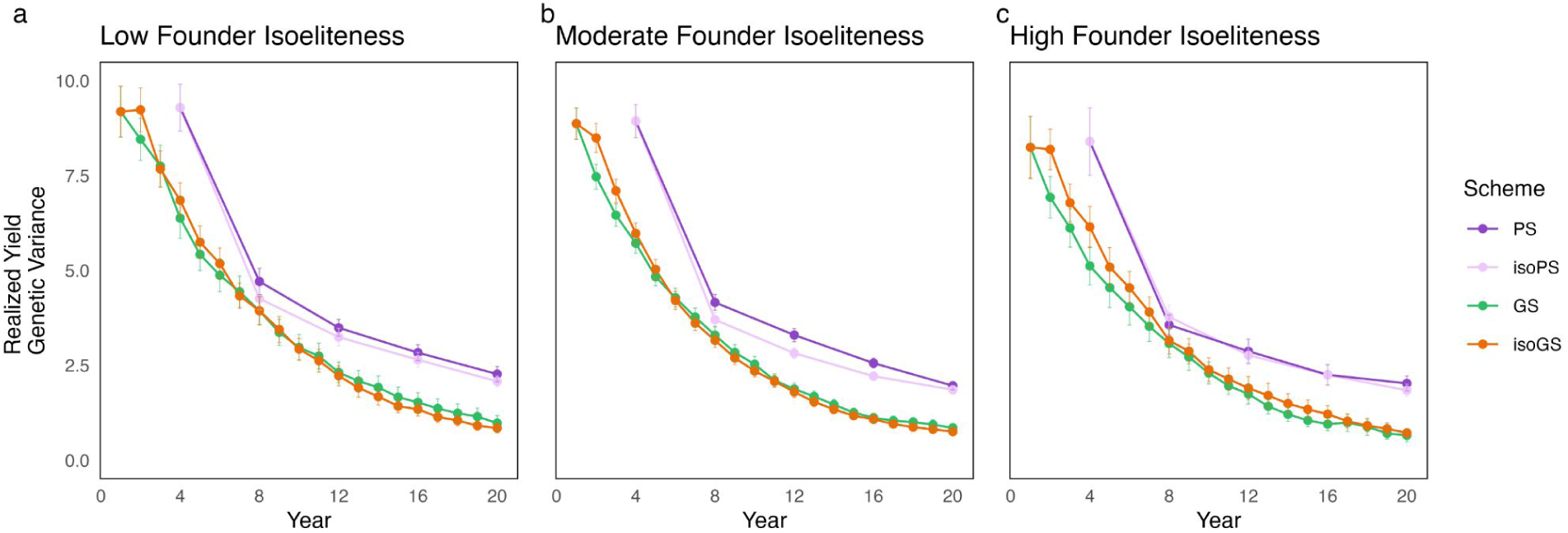
Genetic variance decreased rapidly in GS and isoGS populations. The genetic variance for realized yield was calculated for the F_1_ (S_0_) population at the beginning of each cycle (using the RILs in cycle 0). GS and isoGS populations had a more rapid and sustained decline in variance than PS and isoPS populations. This potentially explains the reduction in GWP accuracy over time (Fig. S3) and the ability of PS to continue driving genetic gain in later cycles while GS plateaued.

**Supplementary Table 1.** Cost estimates for breeding activities.

| **Activity** | **High Labor** | **Low Labor** |
|---|---|---|
| Population establishment + maintenance (1200 plots) | \$15,000.00 | \$1,800.00 |
| Population establishment + maintenance (400 plots) | \$5,000.00 | \$600.00 |
| Population establishment + maintenance (200 plots) | \$3,000.00 | \$400.00 |
| Population establishment + maintenance (100 plots) | \$2,000.00 | \$200.00 |
| Harvest + threshing per plot | \$7.25 | \$1.36 |
| Tissue Collection per plant | \$0.40 | \$0.33 |
| Self per plant | \$0.25 | \$0.19 |
| Cross per plot | \$2.42 | \$1.86 |
| KASP per sample (current) | \$5.00 | \$5.00 |
| KASP per sample (future) | \$1.00 | \$1.00 |
| DArTag per sample (current) | \$10.00 | \$10.00 |
| DArTag per sample (future) | \$2.00 | \$2.00 |

**Supplementary Table 2.** Per-season costs of the first 8 seasons by scheme and labor estimate. Note: Labor costs are shown as “high, low”. “Future” genotyping costs are not shown.

| Season | PS | GS |
|---|---|---|
| 0 | No activities<br>**Total: \$0** | DArTag sequencing for 200 lines: \$5600<br>**Total: \$5600** |
| 1 | Establishment of 200 plots: \$3000, \$400<br>Selfing of 2000 plants: \$500, \$372<br>Harvest of 200 plots: \$1450, \$272<br>**Total: \$4950, \$1044** | Establishment of 100 plots: \$2000, \$200<br>Selfing of 1000 plants: \$250, \$186<br>Harvest of 100 plots: \$725, \$136<br>**Total: \$2975, \$522** |
| 2 | Establishment of 200 plots: \$3000, \$400<br>Selfing of 400 plants: \$100, \$74<br>Harvest of 400 plots: \$290, \$543<br>*isoPS (only for cycle 1):*<br>- *Tissue Collection of 1600 plants: \$635, \$520*<br>- *KASP genotyping of 1600 plants: \$8000*<br>**Total PS: \$5964, \$1018**<br>**Total isoPS: \$14635, \$9538** | Establishment of 100 plots: \$2000, \$200<br>Tissue collection of 800 plants: \$318, \$260<br>DArTag sequencing of 800 plants: \$8000<br>Crossing of 200 plants (closing cycle): \$483, \$372<br>Selfing of 40 plants (for yield trial, YT): \$10, \$7<br>Harvest of 100 + 40 (YT) plots: \$1015, \$190<br>*isoGS (only for cycle 1):*<br>- *KASP genotyping of 800 plants: \$4000*<br>**Total GS: \$11826, \$9030**<br>**Total isoGS: \$15826, \$13030** |
| 3 | Establishment of 400 plots: \$5000, \$600<br>Selfing of 8000 plants: \$2000, \$335<br>Harvest of 400 plots: \$2900, \$543<br>**Total: \$9900, \$2631** | Establishment of 100 + 40 (YT) plots: \$2000, \$200<br>Selfing of 1000 + 800 (YT) plants: \$450, \$335<br>Harvest of 100 + 40 (YT) plots: \$1015, \$190<br>**Total: \$3465, \$725** |
| 4 | Establishment of 400 plots: \$5000, \$600<br>Selfing of 400 plants: \$100, \$74<br>Harvest of 400 plots: \$2900, \$543<br>**Total: \$8000, \$1218** | Establishment of 100 + 120 (YT) plots: \$3000, \$600<br>Tissue collection of 800 plants: \$318, \$260<br>DArTag sequencing of 800 plants: \$8000<br>Crossing of 200 plants (closing cycle): \$483, \$372<br>Selfing of 40 plants (for yield trial, YT): \$10, \$7<br>Yield phenotyping of 120 plots: \$870, \$163<br>Harvest of 100 + 40 (YT) plots: \$1015, \$190<br>**Total: \$13696, \$9393** |
| 5 | Same as season 3<br>**Total: \$9900, \$2631** | Same as season 3<br>**Total: \$3465, \$725** |
| 6 | Establishment of 1200 plots: \$15000, \$1800<br>Selfing of 400 plants: \$100, \$74<br>Yield phenotyping of 1200 plots: \$8700, \$1630<br>**Total: \$23800, \$3504** | Same as season 4<br>**Total: \$13696, \$9393** |
| 7 | Same as season 3<br>**Total: \$9900, \$2631** | Same as season 3<br>**Total: \$3465, \$725** |
| 8 | Establishment of 400 plots: \$5000, \$600<br>Crossing of 200 plants: \$483, \$372<br>Harvest of 400 plots: \$2900, \$543<br>**Total: \$5483, \$972** | Same as season 4<br>**Total: \$13696, \$9393** |
| 9 | Begin cycle 2 | Begin cycle 5 |

**Supplementary Table 3.** Genetic gain across different selection schemes for admixed populations.

| Scheme | Year | Founder Isoeliteness | Mean Gain | Gain Std. Dev. |
|---|---|---|---|---|
| PS | 4 | Low | 9.5 | 1.7 |
| PS | 4 | Moderate | 8.2 | 1.4 |
| PS | 4 | High | 7.0 | 1.1 |
| PS | 12 | Low | 18.7 | 2.7 |
| PS | 12 | Moderate | 17.2 | 2.4 |
| PS | 12 | High | 15.7 | 2.1 |
| PS | 20 | Low | 25.3 | 3.4 |
| PS | 20 | Moderate | 23.4 | 3.1 |
| PS | 20 | High | 21.6 | 2.9 |
| isoPS | 4 | Low | 10.3 | 2.1 |
| isoPS | 4 | Moderate | 8.8 | 1.8 |
| isoPS | 4 | High | 7.0 | 1.4 |
| isoPS | 12 | Low | 19.3 | 2.8 |
| isoPS | 12 | Moderate | 17.6 | 2.5 |
| isoPS | 12 | High | 15.7 | 2.0 |
| isoPS | 20 | Low | 25.7 | 3.5 |
| isoPS | 20 | Moderate | 23.6 | 3.1 |
| isoPS | 20 | High | 21.5 | 2.7 |
| GS | 4 | Low | 8.1 | 4.6 |
| GS | 4 | Moderate | 10.4 | 3.4 |
| GS | 4 | High | 11.1 | 2.4 |
| GS | 12 | Low | 19.5 | 5.0 |
| GS | 12 | Moderate | 19.6 | 4.6 |
| GS | 12 | High | 19.3 | 3.8 |
| GS | 20 | Low | 23.5 | 4.4 |
| GS | 20 | Moderate | 22.2 | 4.6 |
| GS | 20 | High | 21.6 | 4.2 |
| isoGS | 4 | Low | 10.6 | 3.9 |
| isoGS | 4 | Moderate | 11.3 | 2.8 |
| isoGS | 4 | High | 10.3 | 2.1 |
| isoGS | 12 | Low | 21.4 | 4.7 |
| isoGS | 12 | Moderate | 20.5 | 4.1 |
| isoGS | 12 | High | 19.0 | 3.8 |
| isoGS | 20 | Low | 24.8 | 4.5 |
| isoGS | 20 | Moderate | 23.1 | 4.1 |
| isoGS | 20 | High | 21.4 | 4.0 |

## LITERATURE CITED

Allan RE. 1989. Agronomic Comparisons Between Rht1 and Rht2 Semidwarf Genes in Winter Wheat. Crop Sci. 29(5):cropsci1989.0011183X002900050001x. 10.2135/cropsci1989.0011183X002900050001x

Allier A et al. 2020. Optimized breeding strategies to harness genetic resources with different performance levels. BMC Genomics. 21(1):349. 10.1186/s12864-020-6756-0

Anderson TA et al. 2021. Cryptic introgressions contribute to transgressive segregation for early blight resistance in tomato. Theor Appl Genet. 134(8):2561–2575. 10.1007/s00122-021-03842-x

Arends D, Prins P, Jansen RC, Broman KW. 2010. R/qtl: high-throughput multiple QTL mapping. Bioinformatics. 26(23):2990–2992. 10.1093/bioinformatics/btq565

Atlin GN, Cairns JE, Das B. 2017. Rapid breeding and varietal replacement are critical to adaptation of cropping systems in the developing world to climate change. Glob Food Secur. 12:31–37. 10.1016/j.gfs.2017.01.008

Bailey-Serres J et al. 2010. Submergence Tolerant Rice: SUB1’s Journey from Landrace to Modern Cultivar. Rice. 3(2):138–147. 10.1007/s12284-010-9048-5

Barghi N, Hermisson J, Schlötterer C. 2020. Polygenic adaptation: a unifying framework to understand positive selection. Nat Rev Genet. 21(12):769–781. 10.1038/s41576-020-0250-z

Barton NH. 1986. The maintenance of polygenic variation through a balance between mutation and stabilizing selection. Genet Res. 47(3):209–216. 10.1017/S0016672300023156

Barton NH, Etheridge AM, Véber A. 2017. The infinitesimal model: Definition, derivation, and implications. Theor Popul Biol. 118:50–73. 10.1016/j.tpb.2017.06.001

Bernardo R. 1994. Prediction of Maize Single-Cross Performance Using RFLPs and Information from Related Hybrids. Crop Sci. 34(1):cropsci1994.0011183X003400010003x. 10.2135/cropsci1994.0011183X003400010003x

Bernardo R. 2009. Genomewide Selection for Rapid Introgression of Exotic Germplasm in Maize. Crop Sci. 49(2):419–425. 10.2135/cropsci2008.08.0452

Bernardo R. 2025. Why does genomewide prediction become ineffective after several cycles of recurrent selection? Crop Sci. 65(5):e70164. 10.1002/csc2.70164

Bernardo R, Yu J. 2007. Prospects for Genomewide Selection for Quantitative Traits in Maize. Crop Sci. 47(3):1082–1090. 10.2135/cropsci2006.11.0690

Bernardo RN. 2020. Breeding for quantitative traits in plants. Third edition. Stemma Press.

Bouchez A et al. 2002. Marker-Assisted Introgression of Favorable Alleles at Quantitative Trait Loci Between Maize Elite Lines. Genetics. 162(4):1945–1959. 10.1093/genetics/162.4.1945

Boukar O et al. 2020. Introgression Breeding in Cowpea [Vigna unguiculata (L.) Walp.]. Front Plant Sci. 11. 10.3389/fpls.2020.567425

Brodie ED. 2000. Epistasis and the evolutionary process. Wolf JB, Wade MJ, editors. Oxford University Press.

Brown PJ et al. 2011. Distinct Genetic Architectures for Male and Female Inflorescence Traits of Maize. PLOS Genet. 7(11):e1002383. 10.1371/journal.pgen.1002383

Carlborg Ö et al. 2006. Epistasis and the release of genetic variation during long-term selection. Nat Genet. 38(4):418–420. 10.1038/ng1761

Casto AL et al. 2019. Maturity2, a novel regulator of flowering time in Sorghum bicolor, increases expression of SbPRR37 and SbCO in long days delaying flowering. PLOS ONE. 14(4):e0212154. 10.1371/journal.pone.0212154

Cobb JN, Juma RU, et al. 2019. Enhancing the rate of genetic gain in public-sector plant breeding programs: lessons from the breeder’s equation. Theor Appl Genet. 132(3):627–645. 10.1007/s00122-019-03317-0

Cobb JN, Biswas PS, Platten JD. 2019. Back to the future: revisiting MAS as a tool for modern plant breeding. Theor Appl Genet. 132(3):647–667. 10.1007/s00122-018-3266-4

Condón F et al. 2009. Effect of Advanced Cycle Breeding on Genetic Gain and Phenotypic Diversity in Barley Breeding Germplasm. Crop Sci. 49(5):1751–1761. 10.2135/cropsci2008.10.0585

Cooper HD et al., editors. 2001. Broadening the genetic bases of crop production. CABI Pub. in association with Food and Agriculture Organization of the United Nations and International Plant Genetic Resources Institute.

Cooper M, Podlich DW, Smith OS. 2005. Gene-to-phenotype models and complex trait genetics. Aust J Agric Res. 56(9):895. 10.1071/AR05154

Crossa J et al. 2017. Genomic Selection in Plant Breeding: Methods, Models, and Perspectives. Trends Plant Sci. 22(11):961–975. 10.1016/j.tplants.2017.08.011

Daetwyler HD, Hayden MJ, Spangenberg GC, Hayes BJ. 2015. Selection on Optimal Haploid Value Increases Genetic Gain and Preserves More Genetic Diversity Relative to Genomic Selection. Genetics. 200(4):1341–1348. 10.1534/genetics.115.178038

Das B et al. 2024.Strengthening Crop Breeding Networks and Partnerships in Sub Saharan Africa: Results of Three Years of Sustained Effort by CGIAR Accelerated Breeding Initiative.

Dormatey R et al. 2020. Gene Pyramiding for Sustainable Crop Improvement against Biotic and Abiotic Stresses. Agronomy. 10(9):1255. 10.3390/agronomy10091255

Dudley JW. 1982. Theory for Transfer of Alleles. Crop Sci. 22(3):cropsci1982.0011183X002200030049x. 10.2135/cropsci1982.0011183X002200030049x

Edelman GM, Gally JA. 2001. Degeneracy and complexity in biological systems. Proc Natl Acad Sci. 98(24):13763–13768. 10.1073/pnas.231499798

Falconer DS, Mackay T. 2009. Introduction to quantitative genetics. 4. ed., [16. print.]. Pearson, Prentice Hall.

Falk DE. 2010. Generating and maintaining diversity at the elite level in crop breedingThis article is one of a selection of papers from the conference “Exploiting Genome-wide Association in Oilseed Brassicas: a model for genetic improvement of major OECD crops for sustainable farming.” Genome. 53(11):982–991. 10.1139/G10-081

Fall ST et al. 2026. Genomic approaches to build de novo elite breeding gene pools from locally adapted landraces. Theor Appl Genet. 139(1):28. 10.1007/s00122-025-05124-2

Fisher RA. 1919. XV.—The Correlation between Relatives on the Supposition of Mendelian Inheritance. Earth Environ Sci Trans R Soc Edinb. 52(2):399–433. 10.1017/S0080456800012163

Forster BP, Heberle-Bors E, Kasha KJ, Touraev A. 2007. The resurgence of haploids in higher plants. Trends Plant Sci. 12(8):368–375. 10.1016/j.tplants.2007.06.007

Gaynor RC et al. 2017. A Two-Part Strategy for Using Genomic Selection to Develop Inbred Lines. Crop Sci. 57(5):2372–2386. 10.2135/cropsci2016.09.0742

Gaynor RC, Gorjanc G, Hickey JM. 2021. AlphaSimR: an R package for breeding program simulations. G3 GenesGenomesGenetics. 11(2):jkaa017. 10.1093/g3journal/jkaa017

Gholami M et al. 2021. A Comparison of the Adoption of Genomic Selection Across Different Breeding Institutions. Front Plant Sci. 12. 10.3389/fpls.2021.728567

Goddard M. 2009. Genomic selection: prediction of accuracy and maximisation of long term response. Genetica. 136(2):245–257. 10.1007/s10709-008-9308-0

Gorjanc G, Jenko J, Hearne SJ, Hickey JM. 2016. Initiating maize pre-breeding programs using genomic selection to harness polygenic variation from landrace populations. BMC Genomics. 17(1):30. 10.1186/s12864-015-2345-z

Habier D et al. 2010. The impact of genetic relationship information on genomic breeding values in German Holstein cattle. Genet Sel Evol. 42(1):5. 10.1186/1297-9686-42-5

Habier D, Fernando RL, Dekkers JCM. 2007. The Impact of Genetic Relationship Information on Genome-Assisted Breeding Values. Genetics. 177(4):2389–2397. 10.1534/genetics.107.081190

Hallauer AL, Carena MJ. 2012. Recurrent selection methods to improve germplasm in maize. Maydica. 57(4):266–283

Heffner EL, Lorenz AJ, Jannink J-L, Sorrells ME. 2010. Plant Breeding with Genomic Selection: Gain per Unit Time and Cost. Crop Sci. 50(5):1681–1690. 10.2135/cropsci2009.11.0662

Heffner EL, Sorrells ME, Jannink J. 2009. Genomic Selection for Crop Improvement. Crop Sci. 49(1):1–12. 10.2135/cropsci2008.08.0512

Hernandez J, Meints B, Hayes P. 2020. Introgression Breeding in Barley: Perspectives and Case Studies. Front Plant Sci. 11. 10.3389/fpls.2020.00761

Hickey JM, Chiurugwi T, Mackay I, Powell W. 2017. Genomic prediction unifies animal and plant breeding programs to form platforms for biological discovery. Nat Genet. 49(9):1297–1303. 10.1038/ng.3920

Holland JB. 2001. Epistasis and Plant Breeding. In: Janick J, editor. Plant Breeding Reviews. 1st ed. Wiley; p 27–92. 10.1002/9780470650196.ch2

Jannink J-L. 2010. Dynamics of long-term genomic selection. Genet Sel Evol. 42(1):35. 10.1186/1297-9686-42-35

Kannenberg LW, Falk DE. 1995. Models for activation of plant genetic resources for crop breeding programs. Can J Plant Sci. 75(1):45–53. 10.4141/cjps95-008

Kemper KE et al. 2012. Long-term selection strategies for complex traits using high-density genetic markers. J Dairy Sci. 95(8):4646–4656. 10.3168/jds.2011-5289

Kim S. 2015. ppcor: Partial and Semi-Partial (Part) Correlation. https://cran.r-project.org/web/packages/ppcor/index.html

Krchov L-M, Bernardo R. 2015. Relative Efficiency of Genomewide Selection for Testcross Performance of Doubled Haploid Lines in a Maize Breeding Program. Crop Sci. 55(5):2091–2099. 10.2135/cropsci2015.01.0064

Lande R, Thompson R. 1990. Efficiency of Marker-Assisted Selection in the Improvement of Quantitative Traits. Genetics. 124(3):743–756. 10.1093/genetics/124.3.743

Li Xin et al. 2015. Dissecting repulsion linkage in the dwarfing gene Dw3 region for sorghum plant height provides insights into heterosis. Proc Natl Acad Sci. 112(38):11823–11828. 10.1073/pnas.1509229112

Lush JL. 1937. Animal breeding plans.

Mackay IJ, Cockram J, Howell P, Powell W. 2021. Understanding the classics: the unifying concepts of transgressive segregation, inbreeding depression and heterosis and their central relevance for crop breeding. Plant Biotechnol J. 19(1):26–34. 10.1111/pbi.13481

Mackay TFC. 2014. Epistasis and quantitative traits: using model organisms to study gene–gene interactions. Nat Rev Genet. 15(1):22–33. 10.1038/nrg3627

Meuwissen THE, Hayes BJ, Goddard ME. 2001. Prediction of Total Genetic Value Using Genome-Wide Dense Marker Maps. Genetics. 157(4):1819–1829. 10.1093/genetics/157.4.1819

Monyak T, Morris GP. 2026. Emergence of epistatic fitness landscapes from parallel adaptive walks in breeding network metapopulations. G3 GenesGenomesGenetics. jkag306. 10.1093/g3journal/jkag306

Morgante F, Huang W, Maltecca C, Mackay TFC. 2018. Effect of genetic architecture on the prediction accuracy of quantitative traits in samples of unrelated individuals. Heredity. 120(6):500–514. 10.1038/s41437-017-0043-0

Muleta KT et al. 2022. The recent evolutionary rescue of a staple crop depended on over half a century of global germplasm exchange. Sci Adv. 8(6):eabj4633. 10.1126/sciadv.abj4633

Muleta KT, Pressoir G, Morris GP. 2019. Optimizing Genomic Selection for a Sorghum Breeding Program in Haiti: A Simulation Study. G3 GenesGenomesGenetics. 9(2):391–401. 10.1534/g3.118.200932

Olatoye MO et al. 2020. Dissecting Adaptive Traits with Nested Association Mapping: Genetic Architecture of Inflorescence Morphology in Sorghum. G3 GenesGenomesGenetics. 10(5):1785–1796. 10.1534/g3.119.400658

Ortiz-Barrientos D, Jordan D, Cooper M. 2026. The hidden geometry of breeding constraints. Nat Plants. 12(7):1316–1324. 10.1038/s41477-026-02332-6

Paaby AB, Rockman MV. 2014. Cryptic genetic variation: evolution’s hidden substrate. Nat Rev Genet. 15(4):247–258. 10.1038/nrg3688

Podlich DW, Cooper M. 1999. Modelling Plant Breeding Programs as Search Strategies on a Complex Response Surface. In: McKay B et al., editors. Simulated Evolution and Learning. Springer; p 171–178. 10.1007/3-540-48873-1_23

Quinby JR. 1967. The Maturity Genes of Sorghum. In: Norman AG, editor. Advances in Agronomy. Vol. 19. Academic Press; p 267–305. 10.1016/S0065-2113(08)60737-3

R Core Team. 2025. R: a language and environment for statistical computing. R Foundation for Statistical Computing. https://www.R-project.org/

Rasmusson DC, Phillips RL. 1997. Plant Breeding Progress and Genetic Diversity from De Novo Variation and Elevated Epistasis. Crop Sci. 37(2):cropsci1997.0011183X003700020001x. 10.2135/cropsci1997.0011183X003700020001x

Revelle W. 2026. psych: Procedures for Psychological, Psychometric, and Personality Research. https://cran.r-project.org/web/packages/psych/index.html

Rieseberg LH, Archer MA, Wayne RK. 1999. Transgressive segregation, adaptation and speciation. Heredity. 83(4):363–372. 10.1038/sj.hdy.6886170

Rooney WL, Aydin S. 1999. Genetic Control of a Photoperiod-Sensitive Response in Sorghum bicolor (L.) Moench. Crop Sci. 39(2):cropsci1999.0011183X0039000200016x. 10.2135/cropsci1999.0011183X0039000200016x

Rutkoski JE. 2019a. Estimation of Realized Rates of Genetic Gain and Indicators for Breeding Program Assessment. Crop Sci. 59(3):981–993. 10.2135/cropsci2018.09.0537

Rutkoski JE. 2019b. A practical guide to genetic gain. In: Advances in Agronomy. Vol. 157. Academic Press; p 217–249. 10.1016/bs.agron.2019.05.001

Sanchez D et al. 2023. Improving the use of plant genetic resources to sustain breeding programs’ efficiency. Proc Natl Acad Sci. 120(14):e2205780119. 10.1073/pnas.2205780119

Simmonds NW. 1993. Introgression and Incorporation. Strategies for the Use of Crop Genetic Resources. Biol Rev. 68(4):539–562. 10.1111/j.1469-185X.1993.tb01243.x

Swarup S et al. 2021. Genetic diversity is indispensable for plant breeding to improve crops. Crop Sci. 61(2):839–852. 10.1002/csc2.20377

Technow F, Podlich D, Cooper M. 2021. Back to the future: implications of genetic complexity for the structure of hybrid breeding programs. G3 GenesGenomesGenetics. 11(7):jkab153. 10.1093/g3journal/jkab153

Tessele A et al. 2026. Interchromosomal linkage disequilibrium analysis reveals strong indications of sign epistasis in wheat breeding families. G3 GenesGenomesGenetics. 16(1):jkaf265. 10.1093/g3journal/jkaf265

Tibshirani R. 1996. Regression Shrinkage and Selection Via the Lasso. J R Stat Soc Ser B Methodol. 58(1):267–288. 10.1111/j.2517-6161.1996.tb02080.x

Volis S, Shulgina I, Zaretsky M, Koren O. 2011. Epistasis in natural populations of a predominantly selfing plant. Heredity. 106(2):300–309. 10.1038/hdy.2010.79

Watson A et al. 2019. Multivariate Genomic Selection and Potential of Rapid Indirect Selection with Speed Breeding in Spring Wheat. Crop Sci. 59(5):1945–1959. 10.2135/cropsci2018.12.0757

Whitlock MC, Phillips PC, Moore FB-G, Tonsor SJ. 1995. Multiple Fitness Peaks and Epistasis. Annu Rev Ecol Syst. 26:601–629

Wimmer V et al. 2013. Genome-Wide Prediction of Traits with Different Genetic Architecture Through Efficient Variable Selection. Genetics. 195(2):573–587. 10.1534/genetics.113.150078

Windhausen VS et al. 2012. Effectiveness of Genomic Prediction of Maize Hybrid Performance in Different Breeding Populations and Environments. G3 GenesGenomesGenetics. 2(11):1427–1436. 10.1534/g3.112.003699

Wright S. 1935. Evolution in populations in approximate equilibrium. J Genet. 30(2):257–266. 10.1007/BF02982240

Yang CJ et al. 2020. Origin Specific Genomic Selection: A Simple Process To Optimize the Favorable Contribution of Parents to Progeny. G3 GenesGenomesGenetics. 10(7):2445–2455. 10.1534/g3.120.401132

Yang S, Weers BD, Morishige DT, Mullet JE. 2014. CONSTANS is a photoperiod regulated activator of flowering in sorghum. BMC Plant Biol. 14(1):148. 10.1186/1471-2229-14-148

Yu SB et al. 1997. Importance of epistasis as the genetic basis of heterosis in an elite rice hybrid. Proc Natl Acad Sci. 94(17):9226–9231. 10.1073/pnas.94.17.9226

Zebell SG et al. 2025. Cryptic variation fuels plant phenotypic change through hierarchical epistasis. Nature. 1–9. 10.1038/s41586-025-09243-0

